# Single-domain antibodies for targeting, detection and *in vivo* imaging of human CD4^+^ cells

**DOI:** 10.1101/2021.07.02.450848

**Authors:** Bjoern Traenkle, Philipp D. Kaiser, Stefania Pezzana, Jennifer Richardson, Marius Gramlich, Teresa R. Wagner, Dominik Seyfried, Melissa Weldle, Stefanie Holz, Yana Parfyonova, Stefan Nueske, Armin M. Scholz, Anne Zeck, Meike Jakobi, Nicole Schneiderhan-Marra, Martin Schaller, Andreas Maurer, Cécile Gouttefangeas, Manfred Kneilling, Bernd J. Pichler, Dominik Sonanini, Ulrich Rothbauer

## Abstract

The advancement of new immunotherapies necessitates appropriate probes to monitor the presence and distribution of distinct immune cell populations. Considering the key role of CD4^+^ T cells in regulating immunological processes, we generated novel single-domain antibodies (nanobodies, Nbs) that specifically recognize human CD4. After in depth analysis of their binding properties, recognized epitopes, and effects on T cell proliferation, activation and cytokine release, we selected CD4 Nbs that did not interfere with crucial T cell processes *in vitro* and converted them into immune tracers for non-invasive molecular imaging.

By optical imaging, we demonstrate the ability of a high-affinity CD4-Nb to specifically visualize CD4^+^ cells *in vivo* using a xenograft model. Furthermore, time-resolved immune positron emission tomography (immunoPET) of a human CD4 knock-in mouse model showed rapid accumulation of ^64^Cu-radiolabeled CD4-Nb in CD4^+^ T cell-rich tissues. We propose that the CD4 Nbs presented here could serve as versatile probes for stratifying patients and monitoring individual immune responses during personalized immunotherapy in both cancer and inflammatory diseases.

## Introduction

In precision medicine, diagnostic classification of the disease-associated immune status should guide the selection of appropriate therapies. A comprehensive analysis of a patient’s specific immune cell composition, activation state, and infiltration of affected tissue has been shown to be highly informative for patient stratification (Delhalle et al., 2018; Rossi et al., 2019; Scheuenpflug, 2017). CD4^+^ T cells are a key determinant of the immune status due to their essential role in orchestrating immune responses in autoimmune diseases, immune-mediated inflammatory diseases (IMIDs), cancer, and chronic viral infections (Aubert et al., 2011; Becker et al., 1990; Borst et al., 2018; Byrareddy et al., 2016; Chitnis, 2007; Di Mascio et al., 2009; Goverman, 2009; Penaloza-MacMaster et al., 2015). Current diagnostic standards such as intra-cytoplasmic flow cytometry analysis (IC-FACS), immunohistochemistry and ex vivo cytokine assays or RT-PCR analysis are exclusively invasive and limited to endpoint analyses (Doan et al., 2018; Hartmann et al., 2019; Matos et al., 2010; Mousset et al., 2019). Considering the emerging role of infiltrating lymphocytes and the impact of CD4^+^ T cells on the outcome of immunotherapies novel approaches are needed to assess CD4^+^ T cells more holistically (Tay et al., 2021). In this context, non-invasive imaging approaches offer a significant benefit compared to the current diagnostic standard. To date, radiolabeled antibodies have been applied to image CD4^+^ T cells in preclinical models (Di Mascio et al., 2009; Kanwar et al., 2008; Rubin et al., 1996; Steinhoff et al., 2014). Due to the recycling effect mediated by the neonatal Fc receptor, full-length antibodies have a long serum half-life, which requires long clearance times of several days before high-contrast images can be acquired (Dammes and Peer, 2020). Additionally, effector function via the Fc region was shown to induce depletion or functional changes in CD4^+^ cells including the induction of proliferation or cytokine release (Dialynas et al., 1983; Haque et al., 1987; Wilde et al., 1983). Notably, also higher dosages of recombinant antibody fragments like Fab fragments or Cys-diabodies derived from the monoclonal anti-CD4 antibody GK1.5 were recently shown to decrease CD4 expression *in vivo* and inhibit proliferation and IFN-γ production *in vitro* (Freise et al., 2017; Haque et al., 1987; Wilde et al., 1983). These studies highlight the importance of carefully investigating CD4^+^ T cell-specific immunoprobes for their epitopes, binding properties, and functional effects.

During the last decade antibody fragments derived from heavy-chain-only antibodies of camelids (Hamers-Casterman et al., 1993), referred to as VHHs or nanobodies (Nbs) (Hamers-Casterman et al., 1993), have emerged as versatile probes for molecular imaging (reviewed in (Lecocq et al., 2019)). In combination with highly sensitive and/or quantitative whole-body molecular imaging techniques such as optical or radionuclide-based modalities (particularly positron emission tomography (PET)), Nbs have been shown to bind their targets within several minutes of systemic application (Chakravarty et al., 2014). Due to their great potential as highly specific imaging probes, numerous Nbs targeting immune- or tumor-specific cellular antigens are currently in preclinical development and even in clinical trials (Chanier and Chames, 2019; Lecocq et al., 2019; Yang and Shah, 2020).

Here, we generated a set of human CD4-specific Nbs. Following in depth characterization of their binding properties we selected candidates which did not affect T cell proliferation, activation or cytokine release and converted them into immune-tracer for noninvasive optical and PET imaging. Using a mouse xenograft model as well as a human CD4 knock-in mouse model, we successfully demonstrated the capacity of these CD4-Nbs to visualize CD4^+^ cells *in vivo*.

## Results

### Generation of high-affinity CD4 nanobodies

To generate Nbs directed against human CD4 (hCD4), we immunized an alpaca (*Vicugna pacos*) with the recombinant extracellular portion of hCD4 following an 87-day immunization protocol. Subsequently, we generated a Nb phagemid library comprising ∼4 × 10^7^ clones that represent the full repertoire of variable heavy chains of heavy-chain antibodies (VHHs or Nbs) of the animal. We performed phage display using either passively adsorbed purified hCD4 or CHO and HEK293 cells stably expressing full-length human CD4 (CHO-hCD4, HEK293-hCD4 cell lines). Following two cycles of phage display for each condition, we analyzed a total of 612 individual clones by whole-cell phage ELISA and identified 78 positive binders. Sequence analysis revealed 13 unique Nbs representing five different B cell lineages according to their complementarity determining regions (CDR) 3 (Figure 1 A). One representative Nb of each lineage, termed CD4-Nb1 – CD4-Nb5, was expressed in bacteria (*E.coli)* and isolated with high purity using immobilized metal ion affinity chromatography (IMAC) followed by size exclusion chromatography (SEC) (Figure 1 B). To test whether selected Nbs are capable of binding to full-length hCD4 localized at the plasma membrane of mammalian cells, we performed live-cell staining of CHO-hCD4 cells (Figure 1 C, Supplementary Fig. 1). Executed at 4°C, images showed a prominent staining of the plasma membrane, whereas at 37°C the fluorescent signal was mainly localized throughout the cell body, presumably a consequence of endocytotic uptake of receptor-bound Nbs. CHO wt cells were not stained by any of the five CD4-Nbs at both temperatures (data not shown). CD4-Nb1 and CD4-Nb3, both identified by whole-cell panning, displayed strong staining of CHO-hCD4 cells. Of the Nbs derived from panning with recombinant hCD4, CD4-Nb2 also showed strong cellular staining, whereas staining with CD4-Nb4 revealed weak signals. CD4-Nb5 showed no staining under these conditions and was consequently excluded from further analyses (Figure 1 C). To quantitatively assess binding affinities, we performed biolayer interferometry (BLI) measuring serial dilutions of Nbs on the biotinylated extracellular domain of hCD4 immobilized at the sensor tip. For CD4-Nb1 and CD4-Nb2, determined KD values ∼5 and ∼7 nM, respectively, while CD4-Nb3 and CD4-Nb4 displayed lower affinities of 75 nM and 135 nM, respectively (Figure 1 D, Table 1, Supplementary Fig. 2 A). In addition, we determined corresponding EC50 values with full-length plasma membrane-located hCD4 on HEK293-hCD4 cells by flow cytometry. In accordance with cellular staining and biochemically determined affinities these values revealed a strong functional binding for CD4-Nb1 and CD4-Nb2 with EC50 values in the subnanomolar range (∼0.7 nM), whereas CD4-Nb3 and CD4-Nb4 displayed substantially lower cellular affinities (Figure 1 E, Table 1, Supplementary Fig. 2 B). In summary, we generated four CD4-Nbs that bind isolated as well as cell-resident hCD4. While CD4-Nb3 and CD4-Nb4 appeared less affine, CD4-Nb1 and CD4-Nb2 displayed high affinities in the low nanomolar range.

**Figure 1.**
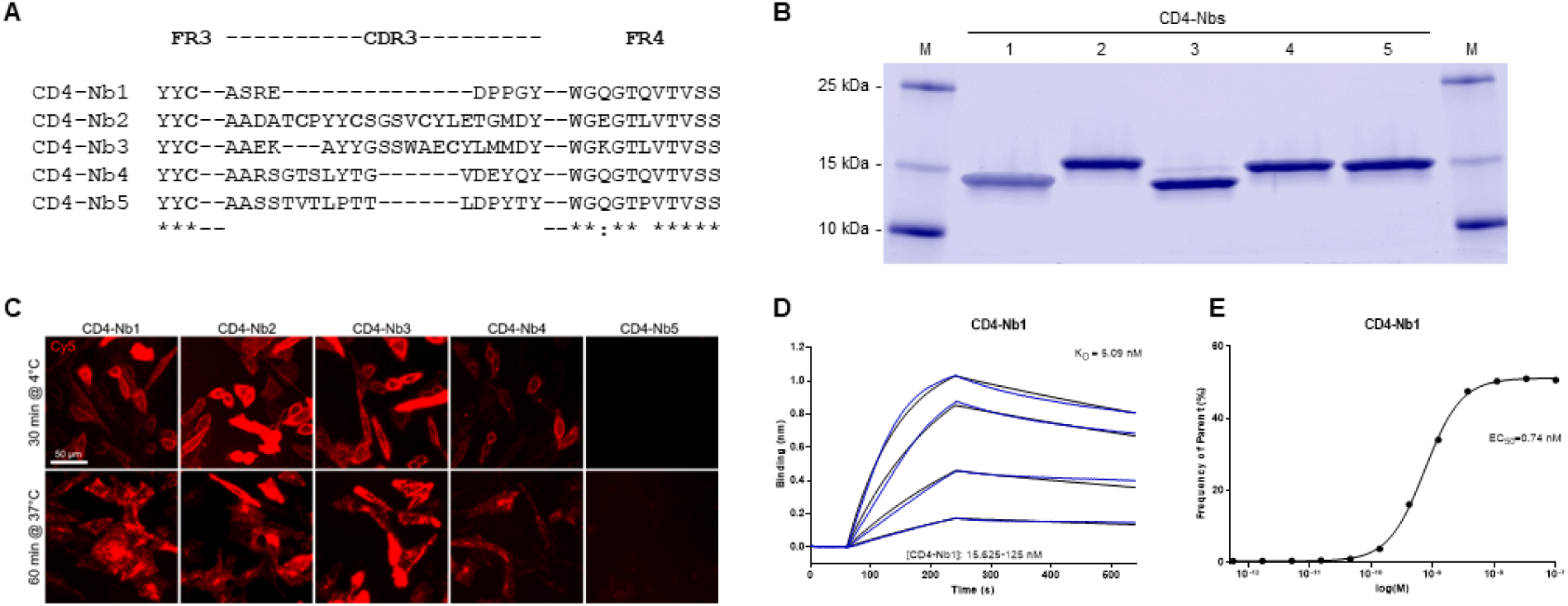
Identification and characterization of Nbs against hCD4. (**A**) Amino acid sequences of the complementarity determining region (CDR) 3 from unique CD4-Nbs selected after two rounds of biopanning are listed. (**B**) Recombinant expression and purification of CD4-Nbs using immobilized metal affinity chromatography (IMAC) and size exclusion chromatography (SEC). Coomassie stained SDS-PAGE of 2 µg of purified Nbs is shown. (**C**) Representative images of live CHO-hCD4 cells stained with CD4-Nbs for 30 min at 4°C (top row) or 60 min at 37°C (bottom row), scale bar: 50 µm. (**D**) For biolayer interferometry (BLI)-based affinity measurements, biotinylated hCD4 was immobilized on streptavidin biosensors. Kinetic measurements were performed using four concentrations of purified Nbs ranging from 15.6 nM - 1000 nM. As an example, the sensogram of CD4-Nb1 at indicated concentrations is shown. (**E**) EC50 determination by flow cytometry. Exemplarily shown for CD4-Nb1, the percentage of positively stained HEK293-hCD4 (frequency of parent) was plotted against indicated concentrations of CD4-Nbs.

**Table 1.**
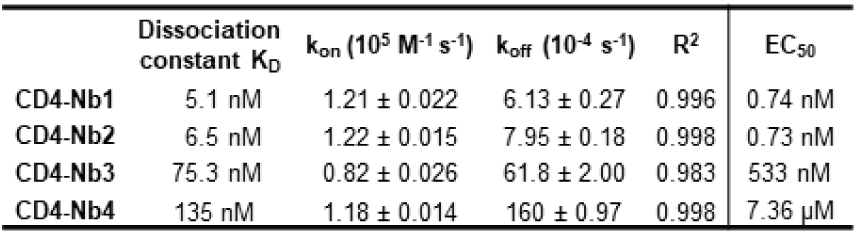
Summary of affinities (KD), association (Kon) and dissociation constants (Koff) determined by BLI (left side) and EC50 values of flow cytometry (right side).

### Domain mapping

Next, we applied chemo-enzymatic coupling using sortase A for site-directed functionalization of CD4-Nbs (Massa et al., 2016; Popp and Ploegh, 2011). We thereby linked peptides conjugated to a single fluorophore to the C-terminus of CD4-Nbs yielding a defined labeling ratio of 1:1 (Virant et al., 2018). Live-cell immunofluorescence imaging showed that all sortase-coupled CD4-Nbs retained their capability of binding to cell-resident hCD4 of CHO-hCD4 cells (Supplementary Fig. 3 A). To localize the binding sites of the selected CD4-Nbs, we generated domain-deletion mutants of hCD4. Expression and correct surface localization of these mutants in CHO cells was confirmed by staining with antibody RPA-T4 binding to domain 1 of CD4. For mutants lacking domain 1, we introduced an N-terminal BC2 tag (Braun et al., 2016) to allow for live-cell surface detection with a fluorescently labeled bivBC2-Nb (Virant et al., 2018) (Supplementary Fig. 3 B). Transiently expressed domain-deletion mutants were then tested for binding of CF568-labeled CD4-Nbs by live-cell immunofluorescence imaging, including a non-specific fluorescently labeled GFP-binding Nb (GFP-Nb) as negative control. Based on these results, we allocated binding of CD4-Nb1 and CD4-Nb3 to domain 1, whereas CD4-Nb2 and CD4-Nb4 bind to domain 3 and/or 4 of hCD4 (Figure 2 A, Supplementary Fig. 3 C).

**Figure 2.**
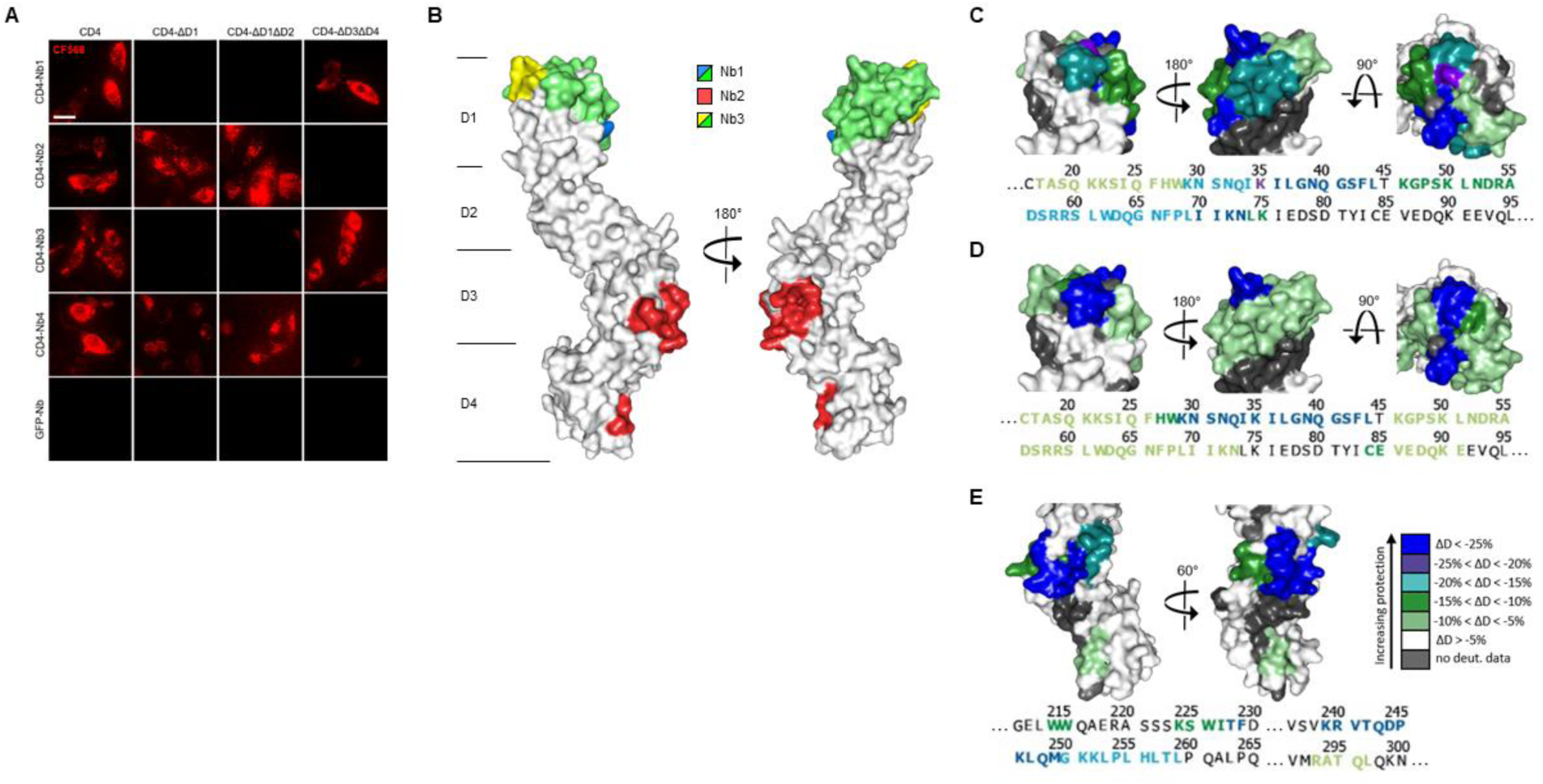
Localization of CD4-Nb binding epitopes. (**A**) Representative images of live CHO cells expressing full-length or domain-deletion mutants of hCD4 stained with fluorescently labeled CD4-Nbs (CF568) are shown, Scale bar 10 µm. (**B**) Surface structure model of hCD4 (PDBe 1wiq) (Wu et al., 1997) and the HDX-MS epitope mapping results of CD4-Nb1 – 3 are depicted. Different colours highlight the amino acid residues protected by CD4-Nb1 (blue), CD4-Nb2 (red) or CD4-Nb 3 (yellow). Overlapping residues protected by both Nb1 and Nb3, are coloured in green. A more detailed surface map (%ΔD) of these specific regions is highlighted in in **C** (CD4-Nb1), **D** (CD4-Nb3) and **E** (CD4-Nb2) with the corresponding CD4 amino acid sequence.

To further examine combinatorial binding of the different CD4-Nbs, we performed an epitope binning analysis by BLI. Recombinant full-length hCD4 was immobilized at the sensor tip and combinations of CD4-Nbs were allowed to bind consecutively (Supplementary Fig. 4). Unsurprisingly, CD4-Nbs binding to different domains, displayed combinatorial binding. Interestingly, a simultaneous binding was also detected for the combination of CD4-Nb1 and CD4-Nb3, suggesting that both CD4-Nbs bind to different epitopes within domain 1. In contrast, we did not observe simultaneous binding for CD4-Nb2 and CD4-Nb4, which might be due to close-by or overlapping epitopes at domain 3/4 for the latter Nb pair.

For a more precise epitope analysis, we conducted a hydrogen-deuterium-exchange (HDX) mass spectrometry analysis of hCD4 bound to CD4-Nb1, CD4-Nb2 or CD4-Nb3 (Figure 2 B-E, Supplementary Fig. 5). Due to its low affinity, CD4-Nb4 was not considered for HDX-MS analysis (data not shown). In accordance with our previous findings, binding of CD4-Nb1 and CD4-Nb3 protected sequences of domain 1 from HDX, whereas CD4-Nb2 protected sequences of domain 3 and 4 of hCD4 (Figure 2 B). The results obtained for binding of CD4-Nb1 (Figure 2 C) are similar to those obtained for CD4-Nb3 (Figure 2 D) in that binding of either Nb reduced hydrogen exchange at amino acid residues (aa) from aa T17 to N73, albeit with a different extent of protection at individual sequence segments. For CD4-Nb1 the greatest protection from HDX was observed for the sequence ranging from aa K35 – L44 corresponding to β strand C’ and C’’ of the immunoglobulin fold of domain 1 and residues aa K46 – K75, comprising β strands D and E. In contrast, binding of CD4-Nb3 confers only a minor reduction in HDX within the latter sequence, but additionally protects sequence aa C84 – E91, which correspond to β strands G and F and their intermediate loop. For CD4-Nb2 we found protection of sequences aa W214 – F229 (β strands C and Ć) and aa K239 – L259 (β strands C’-E), and to a minor extend sequence aa R293 - L296 as part of β strands A of domain 4 (Figure 2 E). In summary, our HDX-MS analysis revealed that all three tested Nbs bind three dimensional epitopes within different parts of hCD4. It further provides an explanation how CD4-Nb1 and CD4-Nb3 can bind simultaneously to domain 1 of hCD4, and confirms that the epitope of CD4-Nb2 is mainly located at domain 3.

### Binding of CD4-Nbs to human PBMCs

Having demonstrated that all selected Nbs bind to recombinant and exogenously overexpressed cellular hCD4, we next examined their capability and specificity of binding to physiologically relevant levels of CD4^+^ T cells within PBMC samples. We co-stained human PBMCs from three donors with CD4-Nbs1-4 coupled to CF568 (100 nM for high-affine CD4-Nb1 and CD4-Nb2; 1000 nM for low-affine CD4-Nb3 and CD4-Nb4) in combination with an anti-CD3 antibody and analyzed the percentage of double-positive cells (CD3^+^CD4^+^) by flow cytometry (Figure 3 A, Supplementary Fig. 6). Compared to staining with an anti-CD4 antibody used as positive control, all CD4-Nbs stained a similar percentage of CD4^+^ T cells for all tested donors, while the non-specific GFP-Nb yielded a negligible percentage of double-positive cells even at the highest concentration (1000 nM) (Figure 3 B, Table 2).

**Figure 3.**
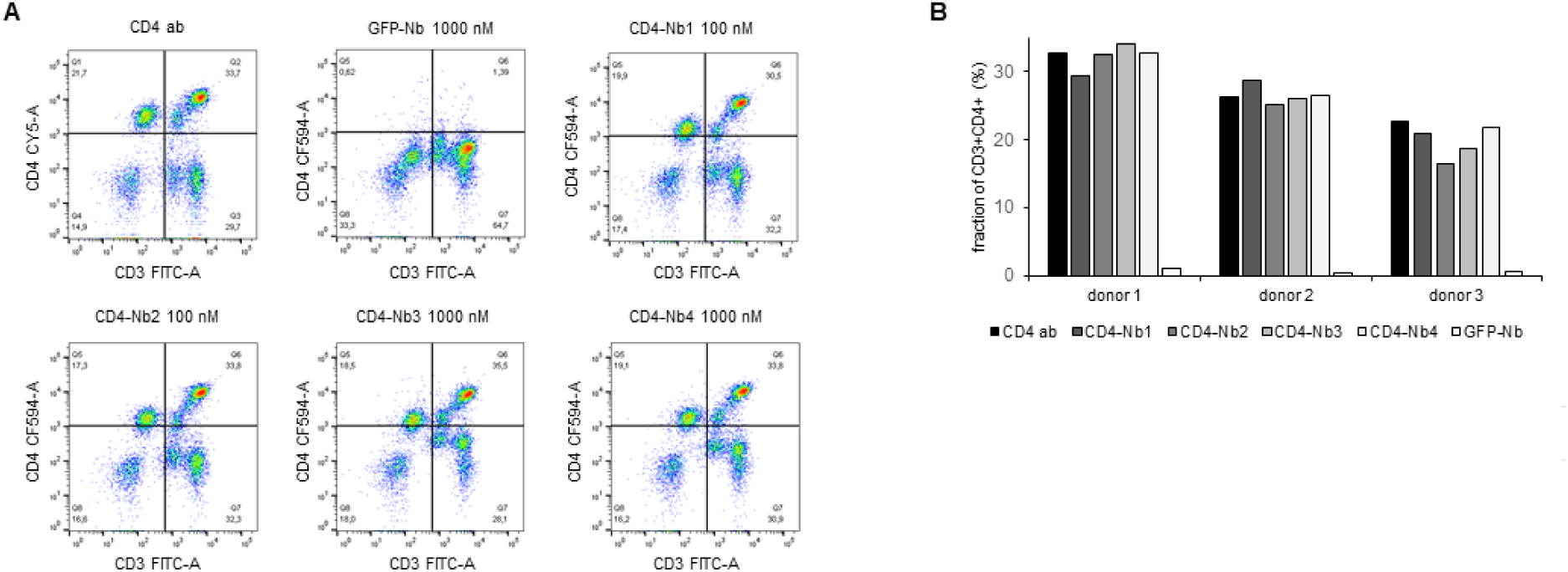
Flow cytometry analysis of human PBMCs stained with fluorescently labeled CD4-Nbs. (**A**) Schematic representation of the final gating step for CD3^+^CD4^+^ double-positive cells derived from donor one. (**B**) Percentage of double-positive cells of three donors, stained with CD4-Nb1 or CD4-Nb2 (100 nM), or CD4-Nb3 or CD4-Nb4 (1000 nM), compared to anti-CD4 antibody and negative control Nb (GFP-Nb, 1000 nM).

**Table 2.**
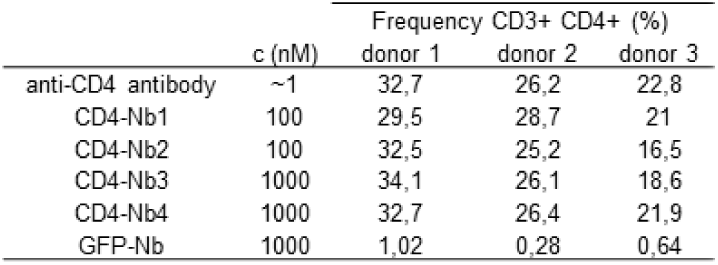
Percentage of double-positive PBMCs from three donors stained with hCD4-Nbs at indicated concentrations.

### Impact of CD4-Nbs on activation, proliferation and cytokine release of CD4^+^ T and immune cells

Towards the envisioned application as clinical imaging tracer, we next assessed basic issues associated with CD4-Nbs, regarding their influence on the activation, proliferation and cytokine release of CD4^+^ T cells. First, to exclude adverse effects of bacterial endotoxins present in the CD4-Nbs1-4 preparations, we attempted to remove endotoxins by depletion chromatography to yield FDA-acceptable endotoxin levels of <0.25 EU per mg. While this was achieved for CD4-Nb1, CD4-Nb4 and the non-specific GFP-Nb, we failed to lower protein-associated endotoxins to acceptable levels for CD4-Nb2 and CD4-Nb3 even after repeated depletion chromatography. Consequently, we continued these experimental studies only with CD4-Nb1, CD4-Nb4 and GFP-Nb as control. In brief, carboxyfluorescein succinimidyl ester (CFSE)-labeled human PBMCs from three pre-selected healthy donors were pre-treated with CD4-Nbs or a control Nb at concentrations between 0.05 µM – 5 µM for 1 h at 37°C, mimicking the expected approximate concentration and serum retention time during clinical *in vivo* imaging application. Cells were then washed to remove Nbs and stimulated with an antigenic (cognate MHCII peptides) or a non-antigenic stimulus (phytohaemagglutinin, PHA-L) and analyzed after 4, 6 and 8 days by flow cytometry with the gating strategy shown in Supplementary Fig. 7 A. According to the highly similar CFSE intensity profiles observed, the total number of cell divisions was not affected by the different Nb treatments (exemplarily shown for one of three donors on day 6, Supplementary Fig. 7 A). For samples of the same donor and time point, no substantial differences in the percentage of proliferated cells were observed between mock incubation and individual Nb treatments.

For both stimuli, the average percentage of proliferated cells increased over time in all donors tested, with no clear differences between conditions (Figure 4 A). As quantitative measure of T cell activation, we also determined the cell surface induction of a very early activation marker (CD69) and of the IL-2 receptor α chain (CD25) on CD4^+^ T cells (Figure 4 B). Among samples of the same donor and stimulation, we found highly similar activation profiles for all Nb treatments. While the percentage of CD4^+^CD25^+^ cells steadily increased over time for MHCII peptide stimulation, for PHA-stimulated condition the percentage of positive cells was similarly high at all times of analysis. Importantly, regardless of the differences between donors, the individual Nb treatments from the same donor did not result in significant differences in the percentage of CD4^+^CD25^+^ or CD4^+^CD69^+^ cells for either stimulation at any point in the analysis.

**Figure 4.**
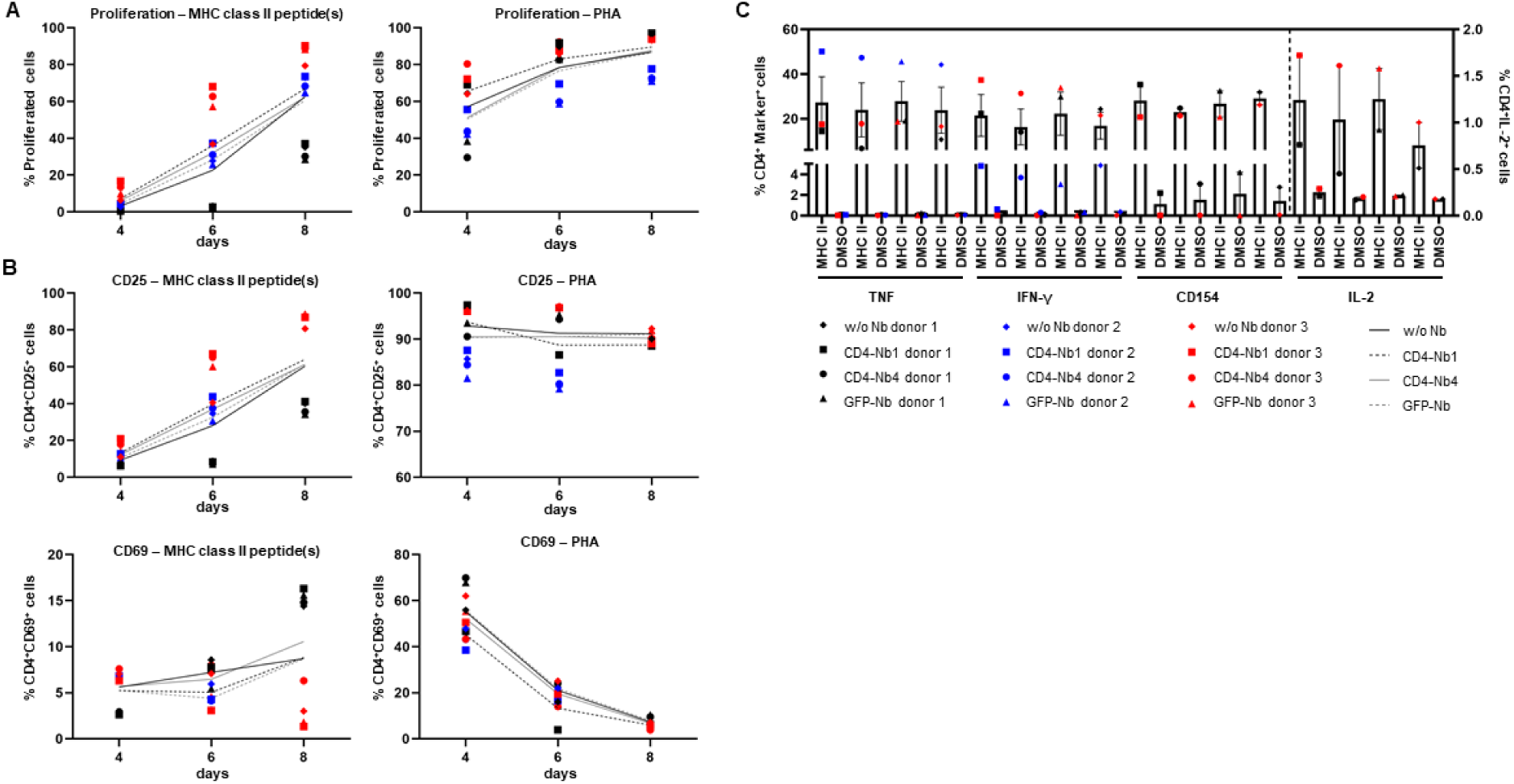
Impact of CD4-Nbs on activation, proliferation and cytokine release of T cells. Cells were stained with CFSE, treated with 5 µM Nbs or without for 1 h (one replicate each), washed and then stimulated with 5 µg/ml MHC-class II peptides, 10 µg/ml PHA or not stimulated, and cultured for 12 days. (**A**) Cells were analyzed by flow cytometry for proliferation (CFSE-low/negative fraction) and activation (CD25 and CD69) on days 4, 6, 8. Proliferation of CD4^+^ cells after stimulation with MHC-class II peptide(s) (left) or PHA (right). (**B**) Activation markers on CD4^+^ cells, Top: CD25 expression after stimulation with MHCII peptide(s) (left) or PHA (right); Bottom: CD69 expression after stimulation with MHCII peptide(s) (left) or PHA (right). Mean percentages of all 3 donors are shown as plain or dotted lines. (**C**) Cytokine and activation marker expression of CD4^+^ cells – TNF, IFN-γ, CD154 (left y-axis) or IL-2 (right y-axis). Cells were restimulated on day 12 with MHC-class II peptide(s) or DMSO (background) for 14 h in the presence of Golgi Stop and Brefeldin A and analyzed by flow cytometry. Error bars display SEM. Gating strategy is shown in **Supplementary** Fig. 6, all percentages are given within CD4^+^ T cells.

Next, we analyzed cytokine expression of CD4^+^ T cells by intracellular cytokine staining after restimulation with cognate MHCII peptides. The corresponding gating strategy is shown in Supplementary Fig. 7 B. Samples of the same donor treated with different Nbs had highly similar percentages of cytokine (TNF, IFN-γ or IL-2) or activation marker (CD154)-positive cells without stimulation and upon stimulation with MHCII peptides (Figure 4 C). Overall, exposure to CD4-Nbs did not affect proliferation, activation or cytokine production of CD4^+^ T cells. In addition, we analyzed potential effects of CD4-Nbs on the release of cytokines from full-blood samples of three further donors. Upon stimulation with lipopolysaccharide (LPS) or PHA-L, we determined the serum concentrations with a panel of pro- and anti-inflammatory cytokines (**Supplementary** Table S2). Although there was significant inter-donor variation for some cytokines, Nb treatment did not result in significant differences in either stimulated or unstimulated samples (Supplementary Fig. 8).

### CD4-Nbs for in vivo imaging

For optical *in vivo* imaging, we labeled CD4-Nbs with the fluorophore Cy5.5 (CD4-Nb-Cy5.5) by sortase-mediated attachment of an azide group followed by click-chemistry addition of DBCO-Cy5.5. First, we tested potential cross-reactivity of the four Cy5.5-labeled CD4-Nbs to murine CD4^+^ lymphocytes. Notably, flow cytometric analysis showed that none of the selected CD4-Nbs bound murine CD4^+^ cells, suggesting exclusive binding to human CD4. Moreover, low-affine binding CD4-Nb4 bound neither mouse nor human CD4^+^ cells at the concentration used here (0.75 µg/ml, ∼49 nM) (Supplementary Fig. 9). Consequently, we focused on CD4-Nb1 as the most promising candidate and CD4-Nb4 as a candidate with a high off-target rate, both of which we further analyzed for their *in vivo* target specificity and dynamic distribution using a murine xenograft model.

To establish human CD4^+^ expressing tumors, NSG mice were inoculated subcutaneously with CD4^+^ T cell leukemia HPB-ALL cells (Masuda et al., 2009). After 2 – 3 weeks, mice bearing HPB-ALL xenografts were *intravenously (i.v.)* injected with 5 µg of CD4-Nb1-Cy5.5, CD4-Nb4-Cy5.5, or a control Nb (GFP-Nb-Cy5.5) and optically imaged in intervals over the course of 24 h (Figure 5 A, Supplementary Fig. 10 A). The Cy5.5 signal intensity (SI) of the control Nb peaked within 10 – 20 minutes and rapidly declined thereafter to approximately the half and a quarter of maximum level at 2 h and 24 h, respectively (Figure 5 B, Supplementary Fig. 10 B). While the SI of the low-affinity CD4-Nb4-Cy5.5 did not exceed the SI of the control Nb at any time (Supplementary Fig. 10 B), CD4-Nb1-Cy5.5 reached its maximum SI within the HPB-ALL xenograft of ∼1.8-fold above the control Nb, at 30 min and slowly declined to ∼90% and ∼80% of maximum after 2 and 4 hours, respectively (Figure 5 B). Based on the differences in the SI between CD4-Nb1-Cy5.5 and GFP-Nb-Cy5.5, we observed constant high target accumulation and specificity between 30 and 480 min post injection (Figure 5 B). After 24 h, mice were euthanized, and the presence of fluorophore-labeled CD4-Nbs within the explanted tumors was analyzed by optical imaging (OI) (Figure 5 C, Supplementary Fig. 10 C). Compared to control, tumors from mice injected with CD4-Nb1-Cy5.5 had ∼4-fold higher Cy5.5 SI, indicating a good signal-to-background ratio for this Nb-derived fluorescently labeled immunoprobe even at later time point. To confirm CD4-specific targeting of CD4-Nb1 within the xenograft, we additionally performed *ex vivo* immunofluorescence of HPB-ALL tumors at 2 h and 24 h post injection (Supplementary Fig. 11). At the early time point, when the *in vivo* OI signal peaked, CD4-Nb1 was widely distributed throughout the whole tumor whereas no Cy5.5 signal was detected in the GFP-Nb-injected mice (Supplementary Fig. 11 A, B). Semiquantitative analysis at the single-cell level revealed intense CD4-Nb1 binding at the surface of HBP-ALL cells that correlated with the CD4 antibody signal and internalization of CD4-Nb1 in some cells (Supplementary Fig. 11 C). In contrast, no binding was observed upon administration of unrelated GFP-Nb (Supplementary Fig. 11 D). 24 h post injection we observed regions of strongly internalized CD4-Nb1 (Supplementary Fig. 11 E, G), but also regions showing a low residual CD4-Nb1 uptake (Supplementary Fig. 11 E, H).

**Figure 5.**
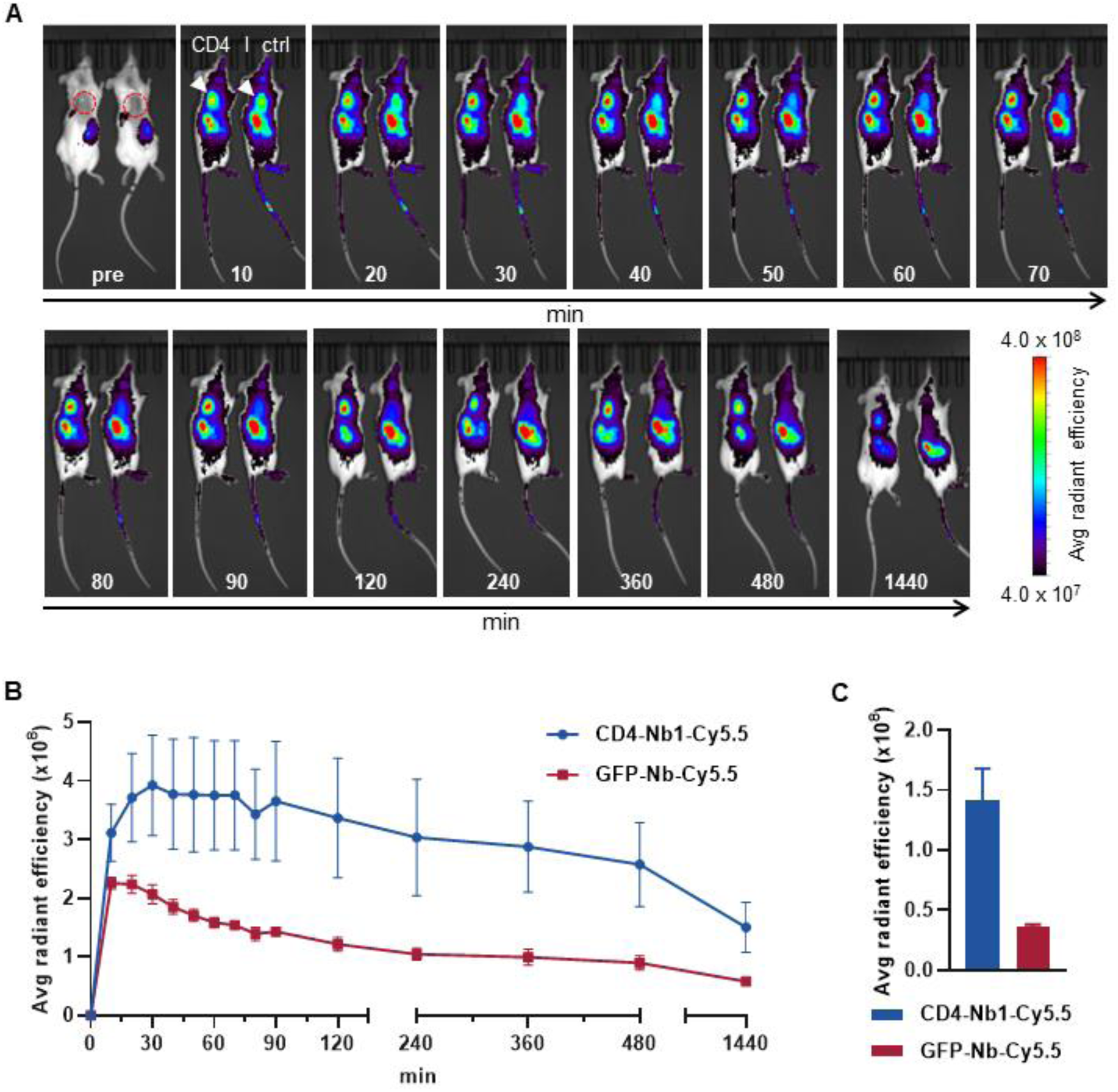
*In vivo* optical imaging (OI) of CD4-Nb1-Cy5.5. 5 µg of CD4-Nb1-Cy5.5 or GFP-Nb-Cy5.5 were administered *i.v.* to *subcutaneously* human CD4^+^ HPB-ALL-bearing NSG mice and tumor biodistribution was monitored by repetitive OI measurements over the course of 24 h. (**A**) Acquired images of each measurement time point of one representative mouse injected with CD4-Nb1-Cy5.5 (left, CD4) or GFP-Nb-Cy5.5 (right, ctrl). Red circles and white arrows indicate the tumor localization at the right upper flank. (**B**) Quantification of the fluorescence signal from the tumors (n = 4 per group, arithmetic mean of the average radiant efficiency ± SEM). (**C**) After the last imaging time point tumors were explanted for *ex vivo* OI, confirming increased accumulation of CD4-Nb1-Cy5.5 compared to the GFP-Nb-Cy5.5 (n = 2 per group, arithmetic mean ± SEM).

The optical imaging data from the xenograft model clearly indicates that the high affinity CD4-Nb1 but not CD4-Nb4 is suitable to specifically visualize CD4^+^ cells *in vivo* within a short period (30 – 120 min) after administration. Considering that this model does not reflect the natural distribution of CD4^+^ T cells in an organism, we continued with a model that allowed us to visualize relevant amounts of CD4^+^ immune cells. Thus we employed a humanized CD4 murine knock-in model (hCD4KI) in which the extracellular fraction of the mouse CD4 antigen was replaced by the human CD4 while normal immunological function and T cell distribution is restored (Killeen et al., 1993).

### ^64^Cu-CD4-Nb1 specifically accumulates in CD4^+^ T cell-rich organs

To generate immunoPET compatible tracers, CD4-Nb1 and GFP-Nb were labeled with the PET isotope ^64^Cu using a copper-chelating BCN-NODAGA group added to our azide-coupled Nbs. Radiolabeling yielded high radiochemical purity (≥95%) and specific binding of ^64^Cu-hCD4-Nb1 to CD4-expressing HBP-ALL cells (46,5 ± 5.6%) *in vitro*, that was ∼30 times higher than the non-specific binding to CD4-negative DHL cells or of the radiolabelled ^64^Cu-GFP-Nb control (Supplementary Fig. 12 A).

Subsequently, we injected ^64^Cu-CD4-Nb1 *i.v.* in hCD4KI and wildtype C57BL/6 mice and performed PET/MR imaging repetitively over 24 h. In two of the hCD4KI animals, we additionally followed tracer biodistribution over the first 90 minutes by dynamic PET (Supplementary Fig. 12 B). As expected for small-sized immunotracers, after an initial uptake peak within the first 10 minutes, ^64^Cu-CD4-Nb1 is rapidly cleared from blood, lung, and liver via renal elimination. In comparison to wildtype, mice carrying the human CD4 antigen on T cells showed an increased tracer accumulation in lymph nodes, thymus, liver, and spleen (Figure 6 A). In these organs, which are known to harbor high numbers of CD4^+^ T cells (Sckisel et al., 2017), we identified 3 h post injection as most suitable imaging time point to discriminate CD4^+^-specific signal from organ background (Figure 6 B). Here, lymph nodes yielded a ∼3-fold, spleen a ∼2.5-fold, and liver a ∼1.4-fold higher ^64^Cu-CD4-Nb1 accumulation in hCD4KI mice compared to wildtype littermates (Figure 6 B). In contrast, we observed similar uptake levels for blood, muscle, lung, and kidney in both groups (Supplementary Fig. 12 C). Analyzing *ex vivo* biodistribution 24 h post tracer injection confirmed persistent accumulation of ^64^Cu-CD4-Nb1 in lymph nodes and spleen of human CD4 expressing mice, although the limited number of animals per group did not allow statistical significance (Supplementary Fig. 12 D). In summary, these results demonstrate that CD4-Nb1 is capable of visualizing and monitoring CD4^+^ T cells in both optical and PET-based imaging.

**Figure 6.**
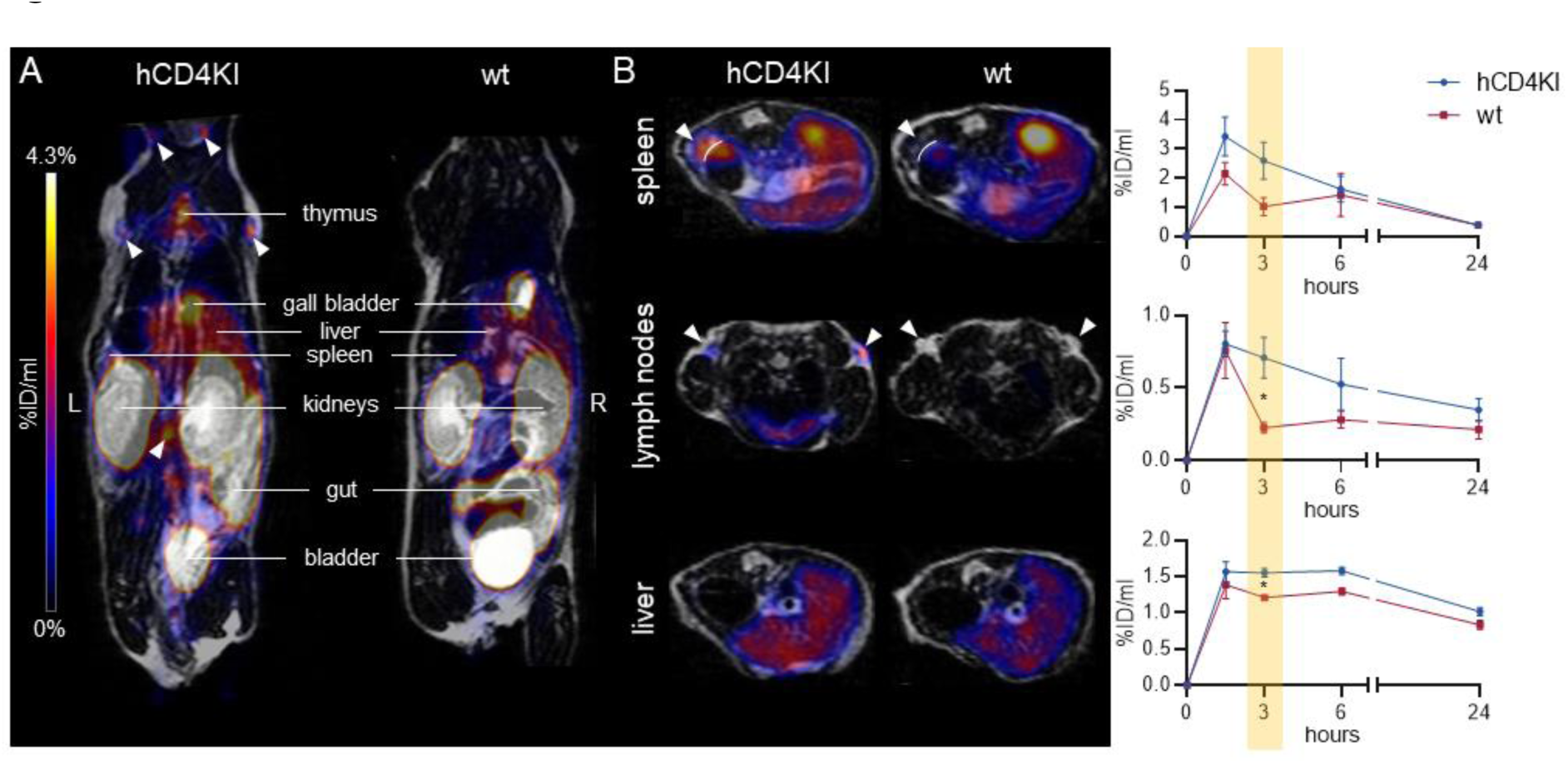
^64^Cu-CD4-Nb1 specifically accumulates in CD4^+^ T cell-rich organs. (**A**) Representative maximum intensity projection PET/MR images of human CD4 knock-in (hCD4KI) and wildtype (wt) C57BL/6 mice 3 h post *i.v.* injection of ^64^Cu-CD4-Nb1. White arrows indicate localization of lymph nodes. (**B**) Exemplary transversal PET/MR images of spleen, lymph nodes, and liver (3 h post injection) and dynamic organ uptake quantification of ^64^Cu-CD4-Nb1 over 24 h (n = 3 per group, arithmetic mean the % injected dose per ml (%ID/ml) ± SEM, unpaired t-test of the 3 h time point, (*) p<0.05). White arrows indicate the target organ.

## Discussion

Given the important role of CD4^+^ T cells, their detailed monitoring is proving to be highly important for the diagnosis and concomitant therapeutic monitoring of a wide variety of diseases. Several mouse studies and early clinical trials have already indicated the value of non-invasive imaging of CD4^+^ cells in rheumatoid arthritis (Steinhoff et al., 2014), colitis (Kanwar et al., 2008), allogenic stem cell transplantation (Tavare et al., 2015), organ transplant rejection (Li et al., 2021), acquired immunodeficiency disease (Di Mascio et al., 2009), and in the context of cancer immunotherapies (Kristensen et al., 2019), using radiolabeled full-length antibodies or fragments thereof. However, biological activity, particularly CD4^+^ T cell depletion, and long-term systemic retention of full-length antibodies limit their development into clinically applied immunoprobes (Choy et al., 2002; Moreland et al., 1995; Steinhoff et al., 2014; Wilde et al., 1983).

The aim of this study was to develop human CD4-specific Nbs as novel *in vivo* imaging probes to overcome the limitations of previous non-invasive imaging approaches. To identify binders that recognize the cellular exposed CD4 we employed two screening strategies where we either selected Nbs against adsorbed recombinant CD4 or against human CD4-expressing cells. Interestingly, both panning strategies proved successful, as demonstrated by the selection of two Nbs each that efficiently bind cell-resident CD4. Combining different biochemical analyses including epitope binning, cellular imaging, and HDX-MS, we were able to elucidate in detail the detected domains, as well as the three-dimensional epitopes addressed by the individual Nbs, and thus identified two candidates, CD4-Nb1 and CD4-Nb3, that can simultaneously bind to different segments within domain 1, while CD4-Nb2 has been shown to bind to domain 3 of CD4. Notably, for most Nbs currently developed for *in vivo* imaging purposes such detailed information is not available (Bala et al., 2018; Blykers et al., 2015; Evazalipour et al., 2014; Huang et al., 2008; Jailkhani et al., 2019; Rashidian et al., 2017; Roovers et al., 2007). However, for CD4-specific Nbs this knowledge is all the more important because epitope-specific targeting of CD4^+^ T cell functions have far-reaching implications. This is true especially for cancer treatment, as CD4^+^ cells have opposing effects on tumor growth and response to immunotherapies, crucially depending on the CD4 effector cell differentiation and tumor entity (Accogli et al., 2021; Bruni et al., 2020). In this context it was shown that domain 1 of CD4 mediates transient interaction of the CD4 receptor and the MHCII complex (Cruikshank et al., 1991; Jonsson et al., 2016; Sakihama et al., 1995), while T cell activation is abrogated when TCR and CD4 colocalization is blocked via domain 3 (Vignali and Vignali, 1999). To further elucidate a possible impact on immunomodulation, we analyzed the effect of CD4-Nb1 and CD4-Nb4 targeting two different domains on CD4^+^ T cell proliferation and cytokine expression. Notably, neither CD4-Nb1 nor CD4-Nb4 affected the behavior of endogenous CD4^+^ T cells *in vitro* or induced increased cytokine levels in whole blood samples when employed at concentrations which are intended for molecular imaging purposes in patients. From these data it can be concluded that these Nbs are mostly biologically inert and thus might be beneficial compared to full-length antibodies (Wilde et al., 1983) or other antibody fragments such as the anti-CD4 Cys-diabody, which was recently reported to inhibit proliferation of CD4^+^ cells and IFN-γ production *in vitro* (Freise et al., 2017).

Following our initial intention to generate immune tracer for *in vivo* imaging, we performed a site-directed labeling approach employing C-terminal sortagging to conjugate an azide group, which can be universally used to attach a multitude of detectable moieties by straightforward DBCO-mediated click chemistry (Rashidian et al., 2017). For the fluorescent and radiolabeled CD4-Nb1, we observed rapid recruitment and sustained staining of CD4^+^ cells in a xenograft and hCD4 knock-in mouse model. Using time resolved PET/MR imaging approach our radiolabeled ^64^Cu-CD4-Nb1 allowed us visualize T cell rich organs with high sensitivity. Beyond lymph nodes we could detect enhanced CD4 specific uptake in liver and spleen. This is a tremendous advantage of an Nb-based tracer, as larger antibody formats tend to accumulate nonspecifically in the spleen and liver due to Fcγ receptor-mediated uptake, making it difficult to distinguish the target-specific signal from background. To further modify serum retention times in order to improve specific tissue targeting, CD4-Nbs could be modified as recently shown by addition of an albumin binding fragment (Tijink et al., 2008) or PEGylation (Rashidian et al., 2017).

In summary, this study demonstrates for the first time the generation and detailed characterization of Nbs specific for human CD4 and their comprehensive experimental evaluation *in vitro* and *in vivo*. In particular, CD4-Nb1 turned out as a promising candidate for non-invasive, whole-body study of CD4^+^ T cells in mice, as well as in humans. Considering the increasing importance of advanced molecular imaging, we anticipate that this Nb-based immunotracer will become a highly versatile tool as a novel theranostic to accompany emerging immunotherapies.

## Data availability

The data that support the findings of this study are available from the corresponding author upon reasonable request.

## Authorship Contributions

B.T., M.K., D.S., B.P. and U.R. designed the study; S.N., A.S. immunized the animal; P.D.K., S.H., Y.P. performed Nb selection; P.D.K, B.T., M.W., T.R.W., A.M. performed biochemical characterization and functionalization of Nbs; M.G., A.Z. performed MS analysis; A.Z. and M.G. performed HDX-MS experiments; J.R., C.G., M.J., N.S.M. analyzed the Nb effects on T cell proliferation and cytokine expression; D.Se., A.M., and S.P. radiolabeled the Nbs; S.P., and D.S. performed *in vivo* imaging; M.S. performed staining of xenograft cryosections; B.T., J.R., C.G., M.K., B.P., D.S. and U.R. drafted the manuscript; M.K., B.P. and U.R. supervised the study.

## Acknowledgements

This work received financial support from the State Ministry of Baden-Wuerttemberg for Economic Affairs, Labour and Tourism (Grant: Predictive diagnostics of immune-associated diseases for personalized medicine. FKZ: 35-4223.10/8). The authors thank Sandra Maier and Ulrich Kratzer (both Natural and Medical Sciences Institute at the University of Tübingen) for technical support with MS analyses, and Birgit Fehrenbacher (Department of Dermatology, University of Tübingen) for technical support with imaging of xenograft cryosections.

## Materials and Methods

### Nanobody library generation

Alpaca immunization and Nb library construction were carried out as described previously (Maier et al., 2015; Traenkle et al., 2015). Animal immunization has been approved by the government of Upper Bavaria (Permit number: 55.2-1-54-2532.0-80-14). In brief, an alpaca (*Vicugna pacos*) was immunized with the purified extracellular domains of human CD4 (aa26-390) recombinantly produced in HEK293 cells (antibodies-online GmbH, Germany). After initial priming with 1 mg, the animal received six boost injections with 0.5 mg hCD4 each, every second week. 87 days after initial immunization, ∼100 ml of blood were collected and lymphocytes were isolated by Ficoll gradient centrifugation using Lymphocyte Separation Medium (PAA Laboratories GmbH). Total RNA was extracted using TRIzol (Life Technologies) and mRNA was transcribed into cDNA using the First-Strand cDNA Synthesis Kit (GE Healthcare). The Nb repertoire was isolated in 3 subsequent PCR reactions using the following primer combinations (1) CALL001 and CALL002, (2) forward primer set FR1-1, FR1-2, FR1-3, FR1-4 and reverse primer CALL002, and (3) forward primer FR1-ext1 and FR1-ext2 and reverse primer set FR4-1, FR4-2, FR4-3, FR4-4, FR4-5 and FR4-6 introducing SfiI and NotI restriction sites. The Nb library was subcloned into the SfiI/ NotI sites of the pHEN4 phagemid vector (Arbabi Ghahroudi et al., 1997).

### Nb screening

For the selection of CD4-specific Nbs two consecutive phage enrichment rounds were performed, both with immobilized recombinant antigen and CHO-hCD4 cells. *E.coli* TG1 cells comprising the hCD4-Nb-library in pHEN4 were infected with the M13K07 helper phage to generate Nb-presenting phages. For each round 1 × 10^11^ phages of the hCD4-Nb-library were applied on immunotubes coated with hCD4 (10 µg/ml). In each selection round extensive blocking of antigen and phages was performed with 5% milk or BSA in PBS-T and with increasing panning rounds PBS-T washing stringency was increased. Bound phages were eluted in 100 mM tri-ethylamine, TEA (pH10.0), followed by immediate neutralization with 1 M Tris/HCl pH7.4. For cell-based panning, 2 × 10^6^ CHO-hCD4 or HEK293-hCD4 were non-enzymatically detached using dissociation buffer (Gibco) and suspended in 5% fetal bovine serum (FBS) in PBS. Antigen expressing cells were incubated with 1 × 10^11^ phages under constant mixing at 4°C for 3 h. Cells were washed 3 × with 5% FBS in PBS. Cell lines were alternated between panning rounds. Phages were eluted with 75 mM citric acid buffer at pH2.3 for 5 min. To deplete non-CD4-specific phages, eluted phages were incubated 3 × with 1 × 10^7^ wt cells. Exponentially growing E.coli TG1 cells were infected with eluted phages from both panning strategies and spread on selection plates for following panning rounds. Antigen-specific enrichment for each round was monitored by counting colony forming unit (CFUs).

### Whole-cell phage ELISA

Polystyrene Costar 96-well cell culture plates (Corning) were coated with poly-L-lysine (Sigma Aldrich) and washed once with H2O. CHO-wt and CHO-hCD4 were plated at 2 × 10^4^ cells per well in 100 µl and grown to confluency overnight. Next day, 70 µl of phage supernatant was added to culture medium of each cell type and incubated at 4°C for 3 h. Cells were washed 5 × with 5% FBS in PBS. M13-HRP-labeled antibody (Progen) was added at a conc. 0.5 ng/ml for 1 h, washed 3 × with 5% FBS in PBS. Onestep ultra TMB 32048 ELISA substrate (Thermo Fisher Scientific) was added and incubated until color change was visible and the reaction was stopped by addition of 100 µl of 1M H2SO4. Detection occurred at 450 nm at a Pherastar plate reader and phage ELISA-positive clones were defined by a 2-fold signal above wt control cells.

### Expression constructs

The cDNA of human CD4 (UniProtKB -P01730) was amplified from hCD4-mOrange plasmid DNA (hCD4-mOrange was a gift from Sergi Padilla Parra; addgene plasmid #110192; http://n2t.net/addgene:110192; RRID:Addgene_110192) by PCR using forward primer hCD4 fwd and reverse primer hCD4 rev and introduced into BamHI and XhoI sites of a pcDNA3.1 vector variant (pcDNA3.1(+)IRES GFP, a gift from Kathleen_L Collins; addgene plasmid #51406; http://n2t.net/addgene:51406; RRID:Addgene_51406). We replaced the neomycin resistance gene (NeoR) with the cDNA for Blasticidin S deaminase (bsd), amplified with forward primer bsd fwd and reverse primer bsd rev, by integration into the XmaI and BssHII sites of the vector. CD4 domain deletion mutants were generated using the Q5 Site-Directed Mutagenesis Kit (NEB) according to the manufactureŕs protocol. For mutants lacking domain 1 of hCD4 we introduced an N-terminal BC2-tag (Braun et al., 2016). For the generation of plasmid pcDNA3.1_CD4_ΔD1_IRES-eGFP we used forward primer ΔD1 fwd and reverse primer ΔD1 rev; for pcDNA3.1_CD4_ΔD1ΔD2_IRES-eGFP forward primer ΔD1ΔD2 fwd and reverse primer ΔD1ΔD2 rev; for pcDNA3.1_CD4_ΔD3ΔD4_IRES-EGFP forward primer ΔD3ΔD4 fwd and reverse primer ΔD3ΔD4 rev. For bacterial expression of Nbs, sequences were cloned into the pHEN6 vector (Rothbauer et al., 2008), thereby adding a C-terminal sortase tag LPETG followed by 6xHis-tag for IMAC purification as described previously (Virant et al., 2018). For protein production of the extracellular domains 1-4 of hCD4 in Expi293 cells, corresponding cDNA was amplified from plasmid DNA containing full-length human CD4 cDNA (addgene plasmid #110192) using forward primer CD4-D1-4 f and reverse primer CD4-D1-4 r. A 6xHis tag was introduced by the reverse primer. Esp3I and EcoRI restriction sites were used to introduce the cDNA into a pcDNA3.4 expression vector with the signal peptide MGWTLVFLFLLSVTAGVHS from the antibody JF5 (Davies et al., 2017).

### Cell culture, transfection, stable cell line generation

HEK293T and CHO-K1 cells were obtained from ATCC (CCL-61, LGC Standards GmbH, Germany). As this study does not include cell line-specific analysis, cells were used without additional authentication. Cells were cultivated according to standard protocols. Briefly, growth media containing DMEM (HEK293) or DMEM/F12 (CHO) (both high glucose, pyruvate, Thermo Fisher Scientific (TFS)) supplemented with 10% (v/v) FBS, L-glutamine and penicillin/streptomycin (P/S; all from TFS) were used for cultivation. Cells were passaged using 0.05% trypsin-EDTA (TFS) and were cultivated at 37°C and 5% CO2 atmosphere in a humidified chamber. Plasmid DNA was transfected using Lipofectamine 2000 (TFS) according to the manufacturer’s protocol. For the generation of the stable HEK293-hCD4 and CHO-hCD4 cell line, 24 h post transfection, cells were subjected to a two-week selection period using 5 µg/ml Blasticidin S (Sigma Aldrich) followed by single cell separation. Individual clones were analyzed by live-cell fluorescence microscopy regarding their level and uniformity of GFP and CD4 expression.

### Protein expression and purification

CD4-specific Nbs were expressed and purified as previously published (Maier et al., 2015; Wagner et al., 2021). Extracellular fragment of human CD4 comprising domains 1-4 of human CD4 and a C-terminal His6-tag was expressed in Expi293 cells according to the manufactureŕs protocol (Thermo Fisher Scientific). Cell supernatant was harvested by centrifugation 4 days after transfection, sterile filtered and purified according to previously described protocols (Becker et al., 2021). For quality control, all purified proteins were analyzed via SDS-PAGE according to standard procedures. Therefore, protein samples were denaturized (5 min, 95°C) in 2x SDS-sample buffer containing 60 mM Tris/HCl, pH6.8; 2% (w/v) SDS; 5% (v/v) 2-mercaptoethanol, 10% (v/v) glycerol, 0.02% bromphenole blue. All proteins were visualized by InstantBlue Coomassie (Expedeon) staining. For immunoblotting proteins were transferred to nitrocellulose membrane (Bio-Rad Laboratories) and detection was performed using anti-His primary antibody (Penta-His Antibody, #34660, Qiagen) followed by donkey-anti-mouse secondary antibody labeled with AlexaFluor647 (Invitrogen) using a Typhoon Trio scanner (GE-Healthcare, excitation 633 nm, emission filter settings 670 nm BP 30).

### Live-cell immunofluorescence

CHO-hCD4, and CHO wt cells transiently expressing CD4 domain-deletion mutants were plated at ∼10,000 cells per well of a µClear 96-well plate (Greiner Bio One, cat. #655090) and cultivated at standard conditions. Next day, medium was replaced by live-cell visualization medium DMEMgfp-2 (Evrogen, cat. #MC102) supplemented with 10% FBS, 2 mM L-glutamine, 2 µg/ml Hoechst33258 (Sigma Aldrich) for nuclear staining, and fluorescently labeled or unlabeled CD4-Nbs at concentrations between 1 nM and 100 nM. Unlabeled CD4-Nbs were visualized by addition of 2.5 µg/ml anti-VHH secondary Cy5 AffiniPure Goat Anti-Alpaca IgG (Jackson Immuno Research). Images were acquired with a MetaXpress Micro XL system (Molecular Devices) at 20× or 40× magnification.

### Biolayer interferometry (BLI)

To determine the binding affinity of purified Nbs to recombinant hCD4, biolayer interferometry (BLItz, ForteBio) was performed. First, CD4 was biotinylated by 3-fold molar excess of biotin-N-hydroxysuccinimide ester. CD4 was then immobilized at single-use streptavidin biosensors (SA) according to manufacturer’s protocols. For each Nb we executed four association/dissociation runs with concentrations appropriate for the affinities of the respective nanobodies (overall between 15.6 nM and 1 µM). As a reference run, PBS was used instead of Nb in the association step. As negative control the GFP-Nb (500 nM) was applied in the binding studies. Recorded sensograms were analyzed using the BLItzPro software and dissociation constants (KD) were calculated based on global fits. For the epitope competition analysis, two consecutive application steps were performed, with a short dissociation period of 30 s after the first association.

### PBMC isolation, cell freezing, and thawing

Fresh blood, buffy coats, or mononuclear blood cell concentrates were obtained from healthy volunteers at the Department of Immunology or from the ZKT Tübingen gGmbH. Participants gave informed consent and the studies were approved by the ethical review committee of the University of Tübingen, projects 156/2012B01 and 713/2018BO2. Blood products were diluted with PBS 1x (homemade from 10x stock solution, Lonza, Switzerland) and PBMCs were isolated by density gradient centrifugation with Biocoll separation solution (Biochrom, Germany). PBMCs were washed twice with PBS 1x, counted with a NC-250 cell counter (Chemometec, Denmark), and resuspended in heat-inactivated (h.i.) fetal bovine serum (Capricorn Scientific, Germany) containing 10% DMSO (Merck). Cells were immediately transferred into a -80°C freezer in a freezing container (Mr. Frosty; Thermo Fisher Scientific). After at least 24 hours, frozen cells were transferred into a liquid nitrogen tank and were kept frozen until use. For the experiments, cells were thawed in IMDM (+L-Glutamin +25mM HEPES; Life Technologies) supplemented with 2.5% h.i. human serum (HS; PanBiotech, Germany), 1x P/S (Sigma-Aldrich), and 50 µm β-Mercaptoethanol (β-ME; Merck), washed once, counted and used for downstream assays.

### Affinity determination by flow cytometry

For cell-based affinity determination, HEK293-hCD4 were detached using enzyme-free cell dissociation buffer (Gibco) and resuspended in FACS buffer (PBS containing 5% FBS). For each staining condition 200,000 cells were incubated with suitable dilution series of CD4 nanobodies at 4°C for 30 min. Cells were washed two times and for detection Cy5 AffiniPure Goat Anti-Alpaca IgG, VHH domain (Jackson ImmunoResearch) was applied for 15 min. PBMCs (Department of Immunology/ ZKT Tübingen gGmbH, Germany) were freshly thawed and resuspended in FACS buffer. For each sample 200,000 cells were incubated with suitable concentrations of CD4 Nbs coupled to CF568 in combination with 1:500 dilution of anti-CD3-FITC (BD Biosciences) at 4°C for 30 min. For control staining PE/Cy5-labeled anti-human CD4 antibody (RPA-T4, Biolegend) was used. After two washing steps, samples were resuspended in 200 µl FACS buffer and analyzed with a BD FACSMelody Cell Sorter. Final data analysis was performed via FlowJo10® software (BD Biosciences).

### Sortase labeling of nanobodies

Sortase A pentamutant (eSrtA) in pET29 was a gift from David Liu (Addgene plasmid # 75144) and was expressed and purified as described (Chen et al., 2011). CF568-coupled peptide H-Gly-Gly-Gly-Doa-Lys-NH2 (sortase substrate) was custom-synthesized by Intavis AG. For the click chemistry a peptide H-Gly-Gly-Gly-propyl-azide was synthesized. In brief, for sortase coupling 50 μM Nb, 250 μM sortase peptide dissolved in sortase buffer (50 mM Tris, pH 7.5, and 150 mM NaCl) and 10 μM sortase were mixed in coupling buffer (sortase buffer with 10 mM CaCl2) and incubated for 4 h at 4°C. Uncoupled Nb and sortase were depleted by IMAC. Unbound excess of unreacted sortase peptide was removed using Zeba Spin Desalting Columns (ThermoFisher Scientific, cat. #89890). Azide-coupled Nbs were labeled by SPAAC (strain-promoted azide-alkyne cycloaddition) click chemistry reaction with 2-fold molar excess of DBCO-Cy5.5 (Jena Bioscience) for 2 h at 25°C. Excess DBCO-Cy5.5 was subsequently removed by dialysis (GeBAflex-tube, 6-8 kDa, Scienova). Finally, to remove untagged Nb, (side product of the sortase reaction), we used hydrophobic interaction chromatography (HIC, HiTrap Butyl-S FF, Cytiva). Binding of DBCO-Cy5.5-coupled Nb occurred in 50 mM H2NaPO4, 1.5 M (NH4)2SO4, pH7.2. Elution took place with 50 mM H2NaPO4, pH7.2. Dye-labeled protein fractions were analyzed by SDS-PAGE followed by fluorescent scanning on a Typhoon Trio (GE-Healthcare, CF568: excitation 532 nm, emission filter settings 580 nm BP 30; Cy5.5 excitation 633 nm, emission filter settings 670 nm BP 30; 546) and subsequent Coomassie staining. Identity and purity of final products were determined by LC-MS (CD4-Nbs-CF568, >60%; CD4-Nb1-Cy5.5, ∼94%; CD4-Nb4-Cy5.5, ∼99%; GBP-Cy5.5; ∼94%, CD4-Nb1-3, ∼99%; bivGFP-Nb, ∼99%).

### Hydrogen-deuterium exchange

#### CD4 deuteration kinetics and epitope elucidation

On basis of the affinity constants of 5.1 nM (CD4-Nb1), 6.5 nM (CD4-Nb2), 75.3 nM (CD4-Nb3) (pre-determined by BLI analysis) the molar ratio of antigen to Nb was calculated ensuring 90% complex formation according to (Kochert et al., 2018). CD4 (5 µL, 65.5 µM) was pre-incubated with CD4-specific Nbs (5 µl; 60.3; 67.4 and 143.1 µM for Nb1; Nb2 and Nb3 respectively) for 10 min at 25°C. Deuteration samples containing CD4 only were pre-incubated with PBS instead of the Nbs. HDX of the pre-incubated samples was initiated by 1:10 dilution with PBS (150 mM NaCl, pH7.4) prepared with D2O leading to a final content of 90% D2O. After 5 and 50 min incubation at 25°C, aliquots of 20 µL were taken and quenched by adding 20 µL ice-cold quenching solution (0.2 M TCEP with 1.5% formic acid and 4 M guanidine HCl in 100 mM ammonium formate solution pH2.2) resulting in a final pH of 2.5. Quenched samples were immediately snap-frozen.

Immobilized pepsin (TFS) was prepared using 60 µl of 50% slurry (in ammonium formate solution pH2.5) that was then dried by centrifugation (1000 x g for 3 min at 0°C) and discarding the supernatant. Prior each analysis, samples were thawed and added to the dried pepsin beads. After digestion for 2 min in a water ice bath the samples were separated by centrifugation at 1000 x g for 30 s at 0°C using a 22 µm filter (Merck, Millipore) and were immediately analyzed by LC-MS. Undeuterated control samples for each complex and CD4 alone were prepared under the same conditions using H2O instead of D2O. Additionally, each Nb was digested without addition of CD4 to generate a list of peptic peptides deriving from the Nb. The HDX experiments of the CD4-Nb-complex were performed in triplicates. The back-exchange of the method as determined using a standard peptide mixture of 14 synthetic peptides was 24%.

#### Chromatography and Mass Spectrometry

HDX samples were analyzed as described previously (Wagner et al., 2021).

### HDX data analysis

A peptic peptide list was generated in a preliminary LC-MS/MS experiment as described previously (Wagner et al., 2021). For data based search no enzyme selectivity was applied, furthermore, identified peptides were manually evaluated to exclude peptides originated through cleavage after arginine, histidine, lysine, proline and the residue after proline (Hamuro and Coales, 2018). Additionally, a separate list of peptides for each nanobody was generated and peptides overlapping in mass, retention time and charge with the antigen digest, were manually removed. Analysis of the deuterated samples was performed in MS mode only and HDExaminer v2.5.0 (Sierra Analytics, USA) was used to calculate the deuterium uptake (centroid mass shift). HDX could be determined for peptides covering 87-88% of the CD4 sequence (Supplementary Fig. 11). The calculated percentage deuterium uptake of each peptide between CD4-Nb and CD4-only were compared. Any peptide with uptake reduction of 5% or greater upon Nb binding was considered as protected. All relevant HDX parameters are shown in Supplementary Table S3 as recommended (Masson et al., 2019).

### Endotoxin determination and removal

The concentration of bacterial endotoxins was determined with Pierce LAL Chromogenic Endotoxin Quantitation Kit (Thermo Fisher Scientific) and removal occurred using EndoTrap HD 1 ml (Lionex) according to the manufacturerś protocols.

### Synthetic peptides

The following HLA-class II peptides were used for the stimulations: MHC class II pool (HCMVA pp65 aa 109-123 MSIYVYALPLKMLNI, HCMV pp65 aa 366-382 HPTFTSQYRIQGKLEYR, EBVB9 EBNA2 aa 276-290 PRSPTVFYNIPPMPL, EBVB9 EBNA1 aa 514-527 KTSLYNLRRGTALA, EBV BXLF2 aa 126-140 LEKQLFYYIGTMLPNTRPHS, EBV BRLF1 aa 119-133 DRFFIQAPSNRVMIP, EBVB9 EBNA3 aa 381-395 PIFIRRLHRLLLMRA, EBVB9 GP350 aa 167-181 STNITAVVRAQGLDV, IABAN HEMA aa 306-318 PKYVKQNTLKLAT,) or CMVpp65 aa 510-524 YQEFFWDANDIYRIF. All peptides were synthesized and dissolved in water 10% DMSO as previously described (purity ≥ 80%), and were kindly provided by S. Stevanović (Loffler et al., 2019).

### Stimulation and cultivation of PBMCs

PBMCs from donors previously screened for *ex vivo* CD4^+^ T cell reactivities against MHC-class II peptides were thawed and rested in T cell medium (TCM; IMDM + 1x P/S + 50μM β-ME + 10% h.i. HS) containing 1μg/mL DNase I (Sigma-Aldrich) at a concentration of 2-3×10^6^ cells/mL for 3h at 37°C and 7.5% CO2. After resting, cells were washed once, counted and up to 1×10^8^ cells were labeled with 1.5-2 μM Carboxyfluorescein succinimidyl ester (CFSE; BioLegend, USA) in 1 mL PBS 1X for 20 min according to the manufacturer’s protocol. The cells were washed twice in medium containing 10% FBS after CFSE labeling and incubated with 5 μM, 0.5 μM, or 0.05 μM of CD4-Nb1, CD4-Nb4 or a control Nb for 1h at 37°C in serum-free IMDM medium. Concentrations and duration were chosen to mimick the expected approximate concentration and serum retention time during clinical application. After incubation, cells were washed twice, counted and each condition was separated into three parts and seeded in a 48-well cell culture plate (1.6-2.5×10^6^ cells/well in triplicates). Cells were stimulated with either 10 μg/ml PHA-L (Sigma-Aldrich), 5 μg/mL MHC class-II peptide(s) or left unstimulated, and cultured at 37°C and 7.5% CO2. 2 ng/mL recombinant human IL-2 (R&D, USA) were added on days 3, 5, and 7. One third of the culture on day 4, one half of the culture on day 6 and day 8, and the remaining cells on day 12 were harvested, counted and stained for flow cytometry analyses. For donor 1, the proliferation/activation status and cytokine production were analyzed in two different experiments, whereas for donors 2 and 3, cells from a single experiment were used for the three assays.

### Assessment of T cell proliferation and activation

Cells from days 4, 6, and 8 were transferred into a 96-well round-bottom plate and washed twice with FACS buffer (PBS + 0.02% sodium azide (Roth, Germany) +2 mM EDTA (Sigma-Aldrich) +2% h.i. FBS). Extracellular staining was performed with CD4 APC-Cy7 (RPA-T4, BD Biosciences), CD8 BV605 (RPA-T8, BioLegend), the dead cell marker Zombie Aqua (BioLegend), CD25 PE-Cy7 (BC96, BioLegend), CD69 PE (FN50, BD Biosciences) and incubated for 20 min at 4°C. All antibodies were used at pre-tested optimal concentrations. Cells were washed twice with FACS buffer. Approx. 500.000 cells were acquired on the same day using a LSRFortessaTM SORP (BD Biosciences, USA) equipped with the DIVA Software (Version 6, BD Biosciences, USA). The percentage of proliferating CD4^+^ cells was determined by assessment of CFSE negative cells, activation by the percentage of CD69^+^ or CD25^+^.

### Assessment of T cell function by intracellular cytokine staining

On day 12, the MHC class II peptide(s)-stimulated and cultured cells were transferred into a 96-well round-bottom plate (0.5 to 1x 10^6^ cells/well) and restimulated using 10 µg/ml of the same peptide(s), 10 µg/ml Staphylococcus enterotoxin B (SEB, Sigma-Aldrich; positive control), or 10% DMSO (negative control). Protein transport inhibitors Brefeldin A (10 µg/ml, Sigma-Aldrich) and Golgi Stop (BD Biosciences) were added at the same time as the stimuli. After 14 h stimulation at 37°C and 7.5% CO2, cells were stained extracellularly with the fluorescently labeled antibodies CD4 APC-Cy7, CD8 BV605, and Zombie Aqua and incubated for 20 min at 4°C. After, cells were washed once, then fixed and permeabilized using the BD Cytofix/Cytoperm solution (BD Biosciences) according to the manufacturer’s instructions, stained intracellularly with TNF Pacific Blue (Mab11), IL-2 PE-Cy7 (MQ1-17H12), IFN-γ AlexaFluor 700 (4S.B7) and CD154 APC (2431) antibodies (all BioLegend) (Widenmeyer et al., 2012) and washed twice. Approx. 500,000 cells were acquired on the same day using a LSRFortessaTM SORP (BD Biosciences, USA), equipped with the DIVA Software (Version 6, BD Biosciences). All flow cytometry analyses were performed with FlowJo version 10.6.2, gating strategies are shown in Supplementary Fig. 6. Statistical analyses were performed with GraphPad Prism version 9.0.0.

### Full blood stimulation and cytokine release assay

100 μl of lithium-heparin blood was incubated for 1 h at 37°C and 7.5% CO2. The blood was stimulated with 5 μM Nb (CD4-Nb1, CD4-Nb4 or control Nb), with 100 ng/mL LPS (Invivogen, USA), or with 2 μg/mL PHA-L in a final volume of 250 μl (serum-free IMDM medium), or left unstimulated for 24 h at 37°C and 7.5% CO2. After two centrifugations, supernatant was collected without transferring erythrocytes. The supernatants were frozen at -80°C until cytokine measurements. Levels of IL-1β, IL-1Ra, IL-4, IL-6, IL-8, IL-10, IL-12p70, IL-13, GM-CSF, IFNγ, MCP-1, MIP-1β, TNFα and VEGF were determined using a set of in house developed Luminex-based sandwich immunoassays each consisting of commercially available capture and detection antibodies and calibrator proteins. All assays were thoroughly validated ahead of the study with respect to accuracy, precision, parallelism, robustness, specificity and sensitivity (EMEA, 2013; FDA, 2018). Samples were diluted at least 1:4 or higher. After incubation of the pre-diluted samples or calibrator protein with the capture coated microspheres, beads were washed and incubated with biotinylated detection antibodies. Streptavidin-phycoerythrin was added after an additional washing step for visualization. For control purposes, calibrators and quality control samples were included on each microtiter plate. All measurements were performed on a Luminex FlexMap® 3D analyzer system, using Luminex xPONENT® 4.2 software (Luminex, USA). For data analysis MasterPlex QT, version 5.0 was employed. Standard curve and quality control samples were evaluated according to internal criteria adapted to the Westgard Rules (Westgard et al., 1981) to ensure proper assay performance.

### Analysis of cross-species reactivity binding to mouse CD4^+^ cells by flow cytometry

Murine CD4^+^ cells were isolated from spleen and lymph nodes of C57BL/6N mice by positive selection over CD4 magnetic microbeads (Miltenyi Biotech, Germany). Human CD4^+^ cells were extracted from blood using StraightFrom® Whole Blood CD4 MicroBeads (Miltenyi Biotech). Single cell suspensions were incubated with 0.75 µg/ml of CD4-Nbs-Cy5.5 (∼47 – 60 nM) or GFP-Nb-Cy5.5 (∼51 nM) in 1% FPS/PBS at 4°C for 20 min and subsequently analyzed on a LSR-II cytometer (BD biosciences).

### Optical Imaging of CD4-expressing HPB-ALL tumors

Human T cell leukemia HPB-ALL cells (German Collection of Microorganisms and Cell Cultures GmbH, DSMZ, Braunschweig, Germany) were cultured in RPMI-1640 supplemented with 10% FBS and 1% P/S. 10^7^ HPB-ALL cells were injected *subcutaneously* in the right upper flank of 7-week-old NOD SCID gamma mice (NSG, NOD.Cg-*Prkdc^scid^ Il2rg^tm1WjI^*/SzJ, Charles River Laboratories, Sulzfeld, Germany) and tumor growth was monitored for 2-3 weeks. When tumors reached a diameter ∼7 mm, 5 µg of CD4-Nbs-Cy5.5 or control Nb (GFP-Nb-Cy5.5) were administered into the tail vein of 2 mice each. Optical imaging (OI) was performed repetitively in short-term isoflurane anesthesia over a period of 24 h using the IVIS Spectrum In Vivo Imaging System (PerkinElmer, Waltham, MA, USA). Four days after the first Nb administration, the CD4-Nbs-Cy5.5 groups received the GFP-Nb-Cy5.5 (and vice versa) and tumor biodistribution was analyzed identically by OI over 24 h. After the last imaging time point, animals were sacrificed and tumors were explanted for *ex vivo* OI analysis. Data were analyzed with Living Image 4.4 software (PerkinElmer). The fluorescence intensities were quantified by drawing regions of interest around the tumor borders and were expressed as average radiant efficiency (photons/s)/(μW/cm^2^) subtracted by the background fluorescence signal before Nb injection to eliminate potential residual signal from the previous Nb application. All mouse experiments were performed according to the German Animal Protection Law and were approved by the local authorities (Regierungspräsidium Tübingen, R5/18).

### Immunofluorescence staining of explanted xenograft tumors

Freshly frozen 5 µm sections of hCD4-Nb1-Cy5.5-containing mice tumors were analyzed using an LSM 800 laser scanning microscope (Zeiss). Afterwards the sections were fixed with perjodate-lysine-paraformaldehyde, blocked using donkey serum and stained with primary rabbit-anti-CD4 antibody (Cell Marque, USA). Bound antibody was visualized using donkey-anti-rabbit-Cy3 secondary antibody (Dianova, Germany). YO-PRO-1 (Invitrogen, USA) was used for nuclear staining. Acquired images of the same areas were manually overlaid.

### Radiolabeling with NODAGA and ^64^Cu

All procedures for conjugation and radiolabeling with ^64^Cu were performed using metal-free equipment and Chelex 100 (Sigma-Aldrich) pretreated buffers. Azide-modified Nbs (100 µg) were treated with 4 µl of 5 mM EDTA in 250 mM sodium acetate buffer (pH 6) for 30 min at RT. The protein was reacted with 15 molar equivalents of BCN-NODAGA (CheMatech, Dijon France) in 250 mM sodium acetate pH 6 for 30 min at RT followed by incubation at 4 °C for 18 h. Excess of chelator was removed by ultrafiltration (Amicon Ultra 0.5 ml, 3 kDa MWCO, Merck Millipore) using the same buffer. [^64^Cu]CuCl2 (150 MBq in 0.1 M HCl) was neutralized by addition of 1.5 volumes of 0.5 M ammonium acetate solution (pH 6), resulting in a pH of 5.5. To this solution, 50 µg of conjugate was added and incubated at 42 °C for 60 min. 1 µl of 20% DTPA solution was added to quench the labeling reaction. Complete incorporation of the radioisotope was confirmed after each radiosynthesis by thin-layer chromatography (iTLC-SA, Agilent Technologies; mobile phase 0.1 M citric acid pH 5) and high-performance size exclusion chromatography (HPSEC, BioSep SEC-s2000, 300 x 7.8 mm, Phenomenex; mobile phase DPBS with 0.5 mM EDTA). All radiolabeled preparations used for *in vivo* PET imaging had radiochemical purities of ≥97% (iTLC) and ≥94% (HPSEC).

### In vitro radioimmunoassay

10^7^ HPB-ALL or DHL cells were incubated in triplicates with 1 ng (3 MBq/µg) of radiolabeled ^64^Cu-CD4-Nb1 or ^64^Cu-GFP-Nb for 1 h at 37°C and washed twice with PBS/2% FCS. The remaining cell-bound radioactivity was measured using a Wizard² 2480 gamma counter (PerkinElmer Inc., Waltham, MA, USA) and quantified as percentage of total added activity.

### PET/MR imaging

Human CD4 knock-in (hCD4KI, genOway, Lyon, France) and wildtype C57BL/6J mice (Charles River) were injected intravenously with 5 µg (∼15 MBq) of ^64^Cu-CD4-Nb1. During the scans, mice were anesthetized with 1.5% isoflurane in 100% oxygen and warmed by water-filled heating mats. Ten-minute static PET scans were performed after 1.5, 3, 6, and 24 h in a dedicated small-animal Inveon microPET scanner (Siemens Healthineers, Knoxville, Tennessee, USA; acquisition time: 600 s). For anatomical information, sequential T2 TurboRARE MR images were acquired immediately after the PET scans on a small animal 7 T ClinScan magnetic resonance scanner (Bruker BioSpin GmbH, Rheinstetten, Germany). Dynamic PET data of the first 90 minutes post injection were gained in two mice and divided into 10-minute-frames. After attenuation correction by a cobalt-57 point source, PET images were reconstructed using an ordered subset expectation maximization (OSEM3D) algorithm and analyzed with Inveon Research Workplace (Siemens Preclinical Solutions). The volumes of interest of each organ were drawn based on the anatomical MRI to acquire corresponding PET tracer uptake. The resulting values were decay-corrected and presented as percentage of injected dose per volume (%ID/ml). *Ex vivo* γ-counting was conducted after the last imaging time point by measuring the weight and radioactivity of each organ. For quantification, standardized aliquots of the injected radiotracer were added to the measurement.

### Analyses and Statistics

Data analysis of the flow cytometry data was performed with the FlowJo Software Version 10.6.2 (FlowJo LLT, USA) and graph preparation and statistical analysis was performed using the GraphPad Prism Software (Version 8.3.0 or higher).

## Supplementary Information

**Supplementary Figure 1.**
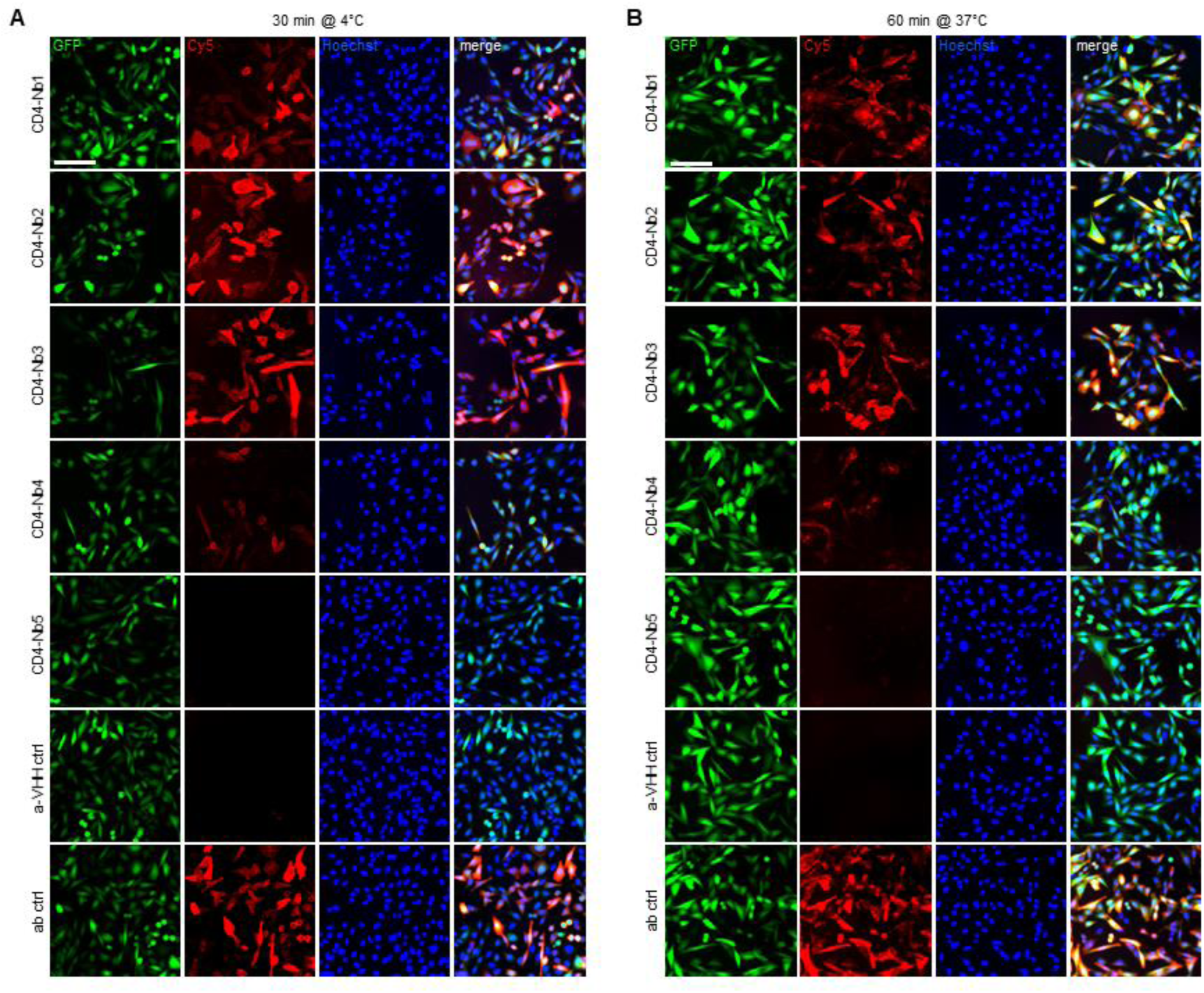
Live-cell immunofluorescence staining of CHO-hCD4 cells incubated with CD4-Nbs (100 nM) followed by detection using a secondary Cy5-labeled anti-VHH antibody and 2 µg/ml Hoechst33258 for 30 min at 4°C (**A**) or 60 min at 37°C (**B**). Shown are representative images of CHO-hCD4 cells simultaneously expressing cytosolic GFP (left column) and hCD4 (second column from left) from a bicistronic mRNA. Nuclear staining and merge of channels is shown in column 3 and 4 from the left. Negative control staining using secondary Cy5-labeled anti-VHH antibody alone (a-VHH ctrl) and positive control using Cy5-labeled anti-hCD4 antibody (ab ctrl) are shown in bottom two rows. Scale bar 100 µm.

**Supplementary Figure 2.**
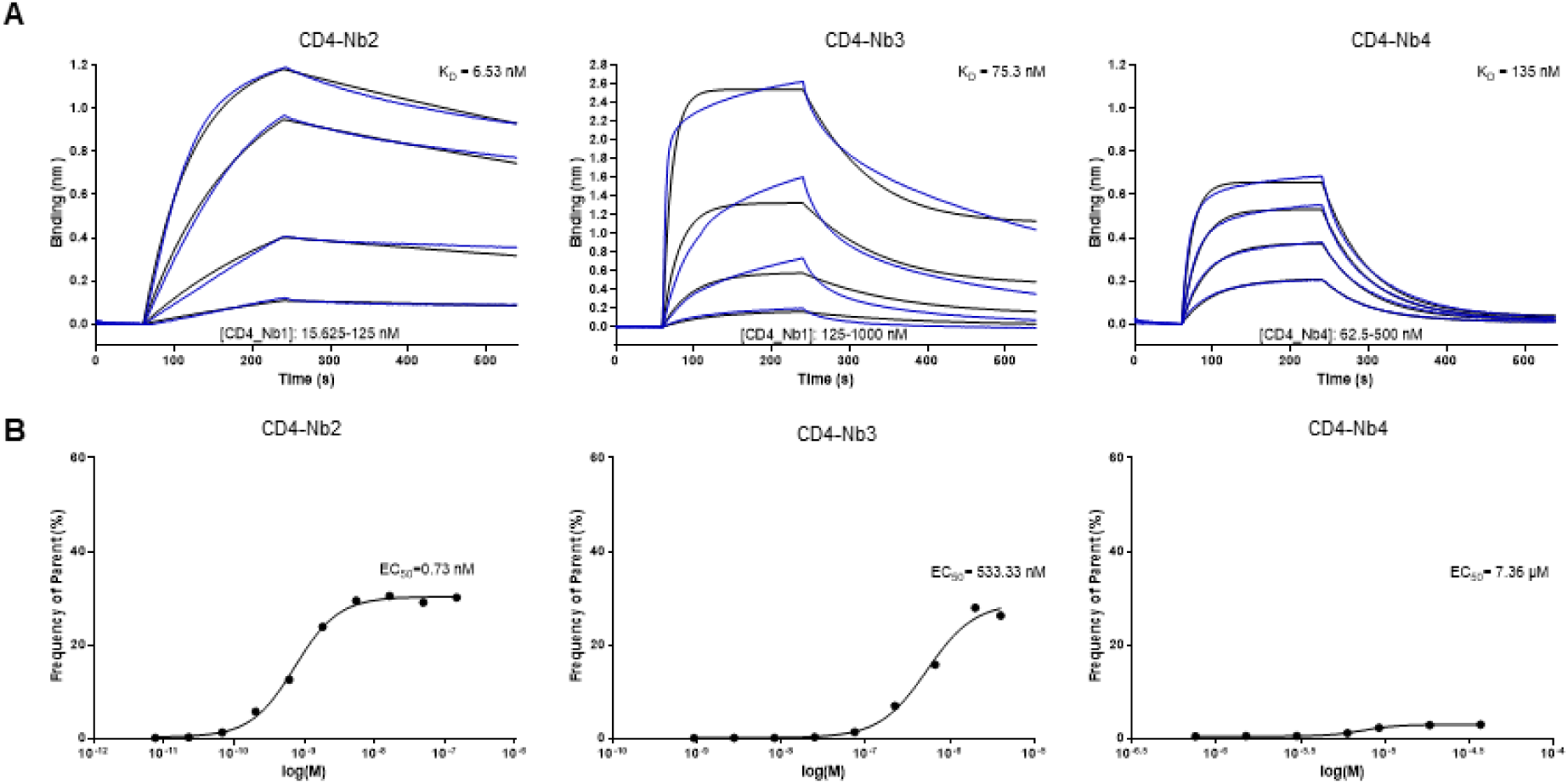
Affinities of identified CD4-Nbs (**A**) Sensograms of biolayer interferometry-based affinity measurements of CD4-Nb2, CD4-Nb3 and CD4-Nb4 are shown. Biotinylated hCD4 was immobilized on streptavidin biosensors and kinetic measurements were performed by using four concentrations of purified Nbs ranging from 15.6 nM -1 µM. (**B**) EC50 determination of CD4-Nbs for cellularly expressed hCD4 by flow cytometry. The percentage of positively stained HEK293-hCD4 (frequency of parent) was plotted against indicated concentrations of CD4-Nbs. EC50 values were calculated from a four-parametric sigmoidal model.

**Supplementary Figure 3.**
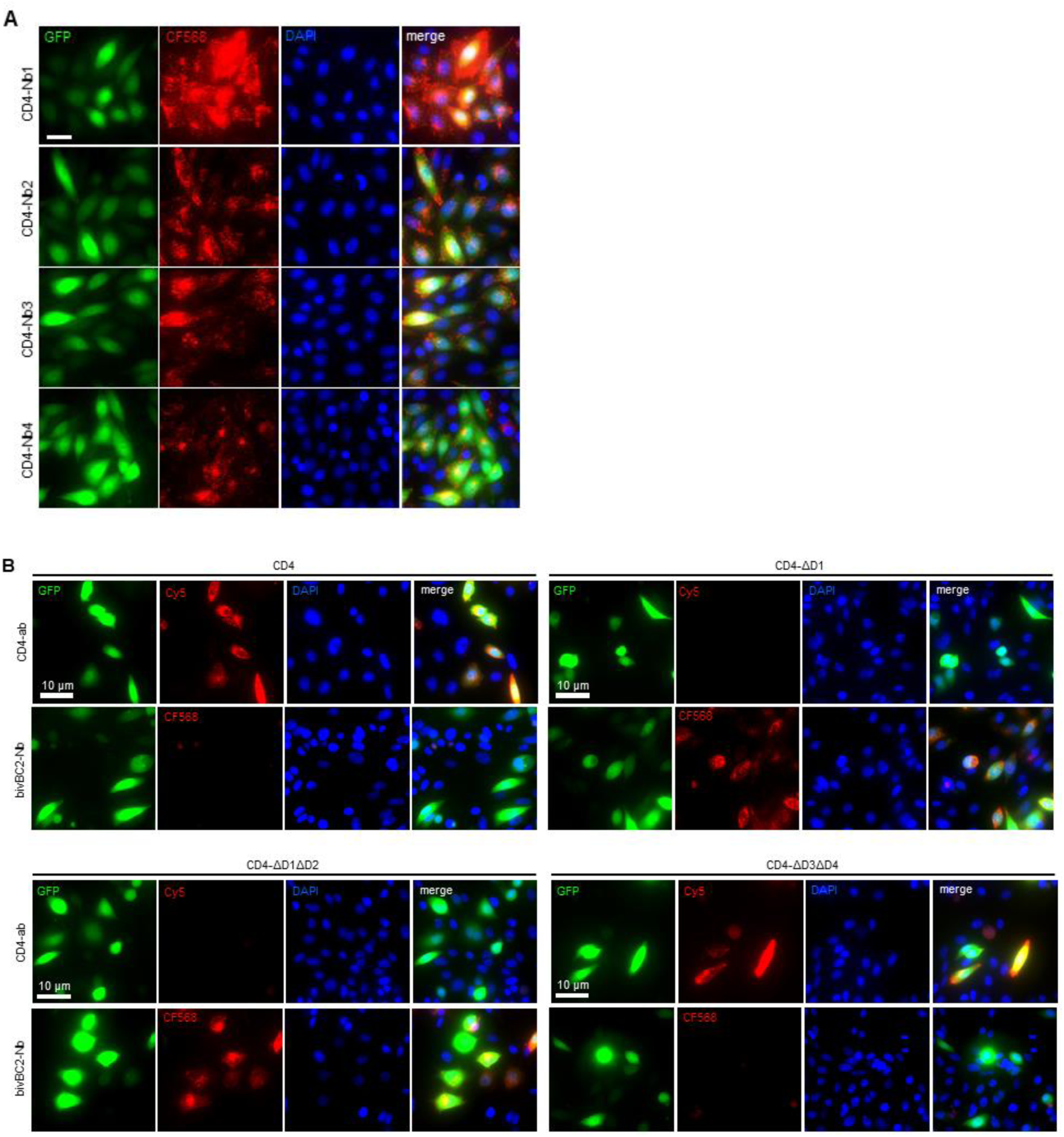
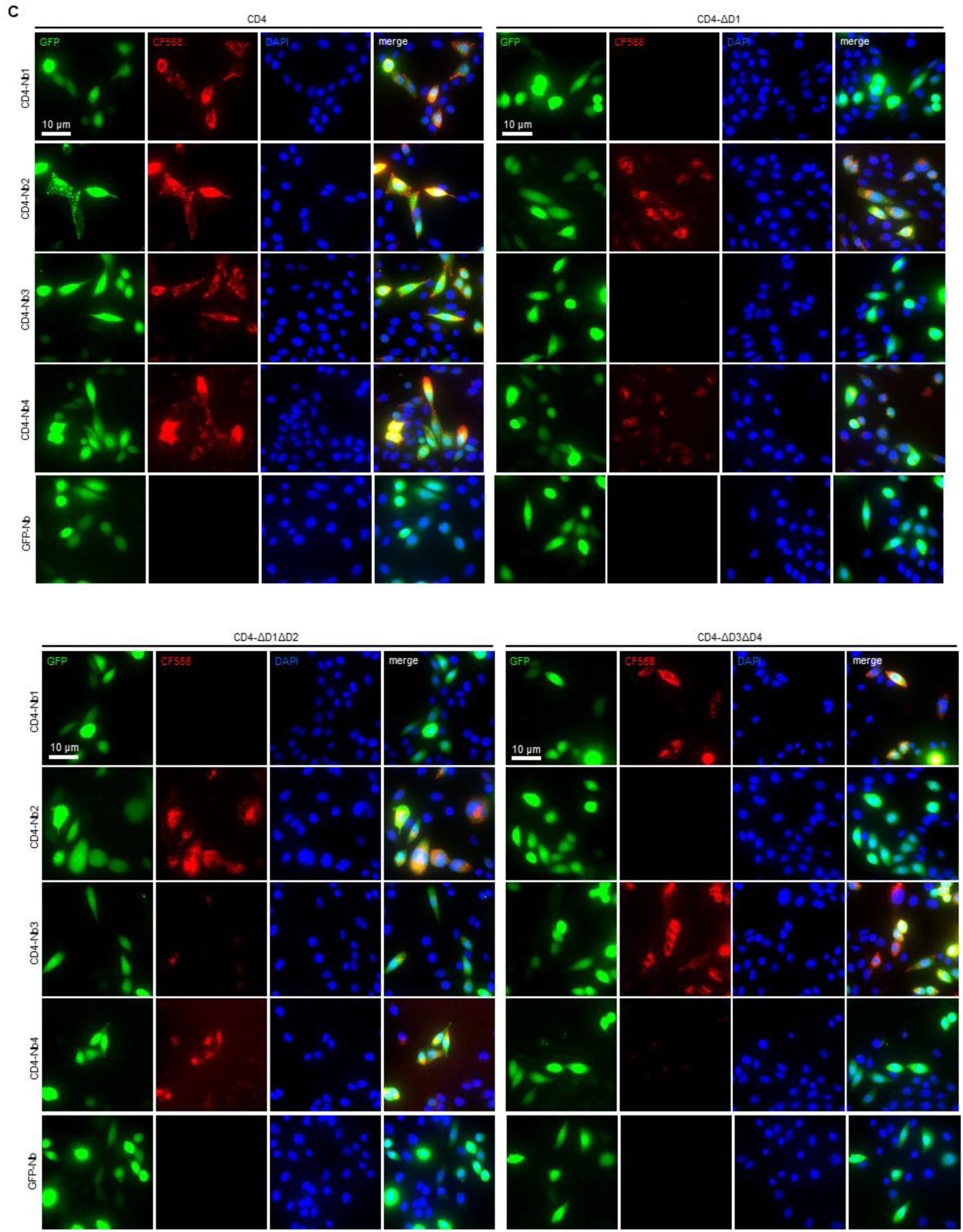
CD4-Nbs bind different domains of human CD4. (**A**) Live-cell immunofluorescence staining of CHO-hCD4 cells with CD4-Nbs coupled to the fluorescent dye CF568, and 2 µg/ml Hoechst33258 for 60 min at 37°C. (**B**) Control staining of full-length hCD4 (CD4) or hCD4 domain-deletion mutants CD4-ΔD1, CD4-ΔD1ΔD2 or CD4-ΔD3ΔD4 with fluorescently labeled anti-CD4 antibody RPA-T4-PE/Cy5 (CD4-ab) or bivalent BC2-Nb coupled to CF568 (bivBC2-Nb). (**C**) Live-cell immunofluorescence staining of CHO cells transiently expressing full-length hCD4 or hCD4 domain-deletion mutants with CF568-labeled CD4-Nbs or a non-specific GFP-Nb (100 nM). Scale bars 10 µm.

**Supplementary Figure 4.**
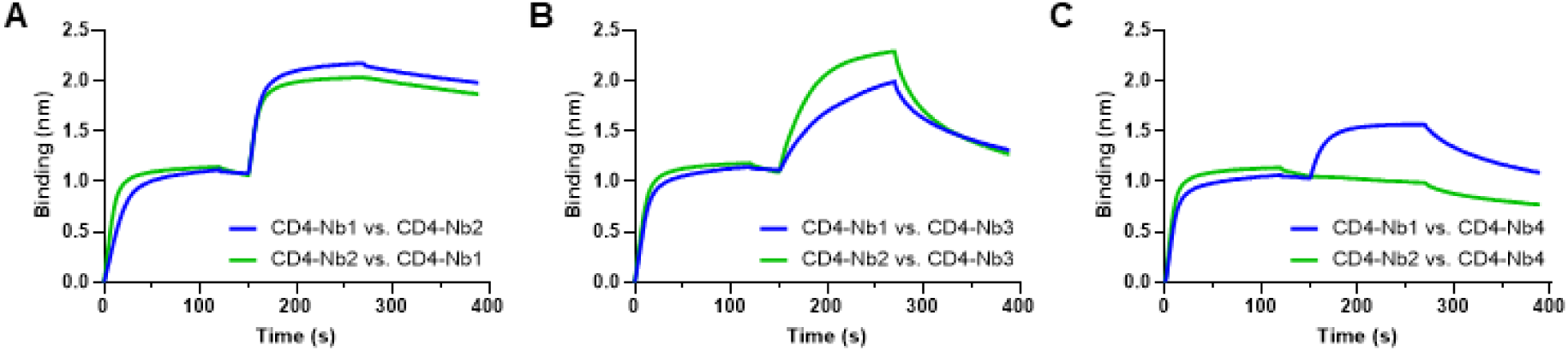
Epitope binning analysis of CD4-Nbs by biolayer interferometry (BLI) (**A**) Representative BLI sensograms of single measurements of combinatorial Nb binding to the recombinant extracellular portion of hCD4 of CD4-Nb1 (blue) and CD4-Nb2 (green) with (**A**) one another, (**B**) CD4-Nb3, or (**C**) CD4-Nb4.

**Supplementary Figure 5.**
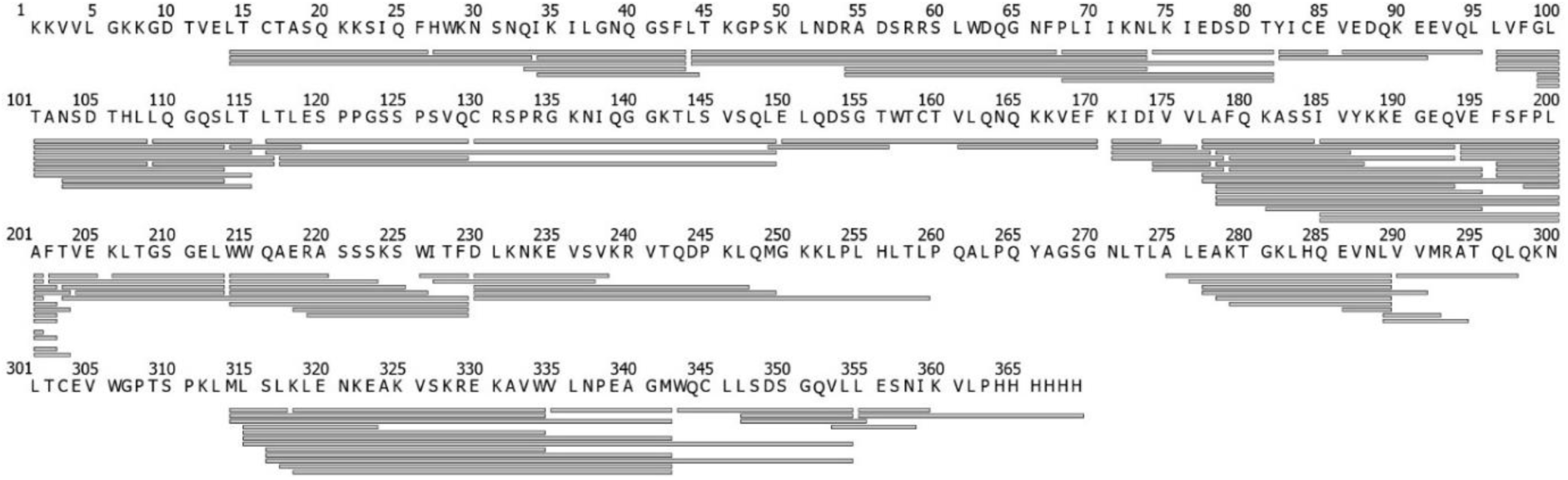
Peptide sequence coverage of human CD4 for HDX-MS analysis. 116 possible peptides could be identified by MSMS (depicted as bars) leading to a sequence coverage of 88% for the HDX analysis.

**Supplementary Figure 6.**
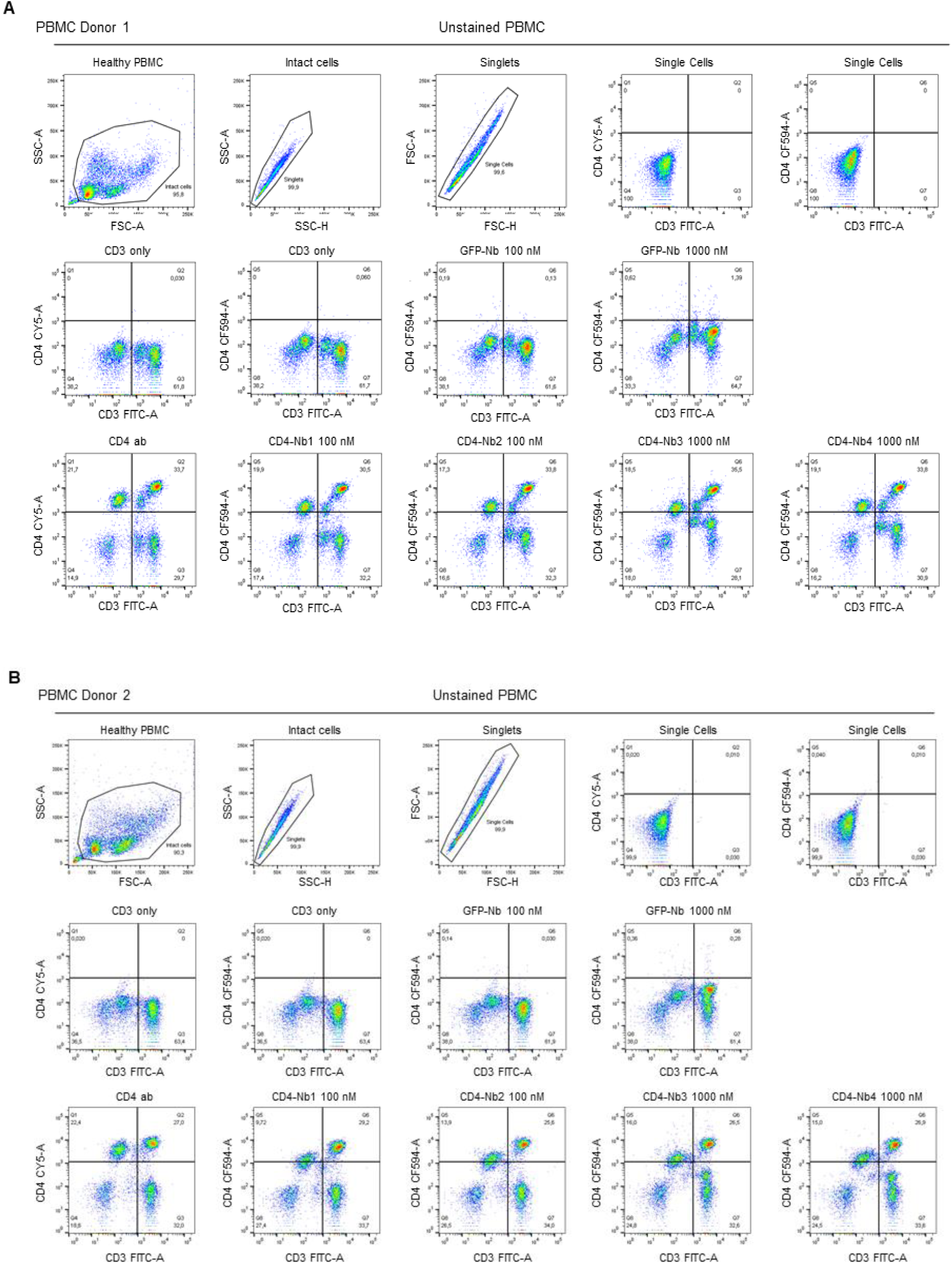
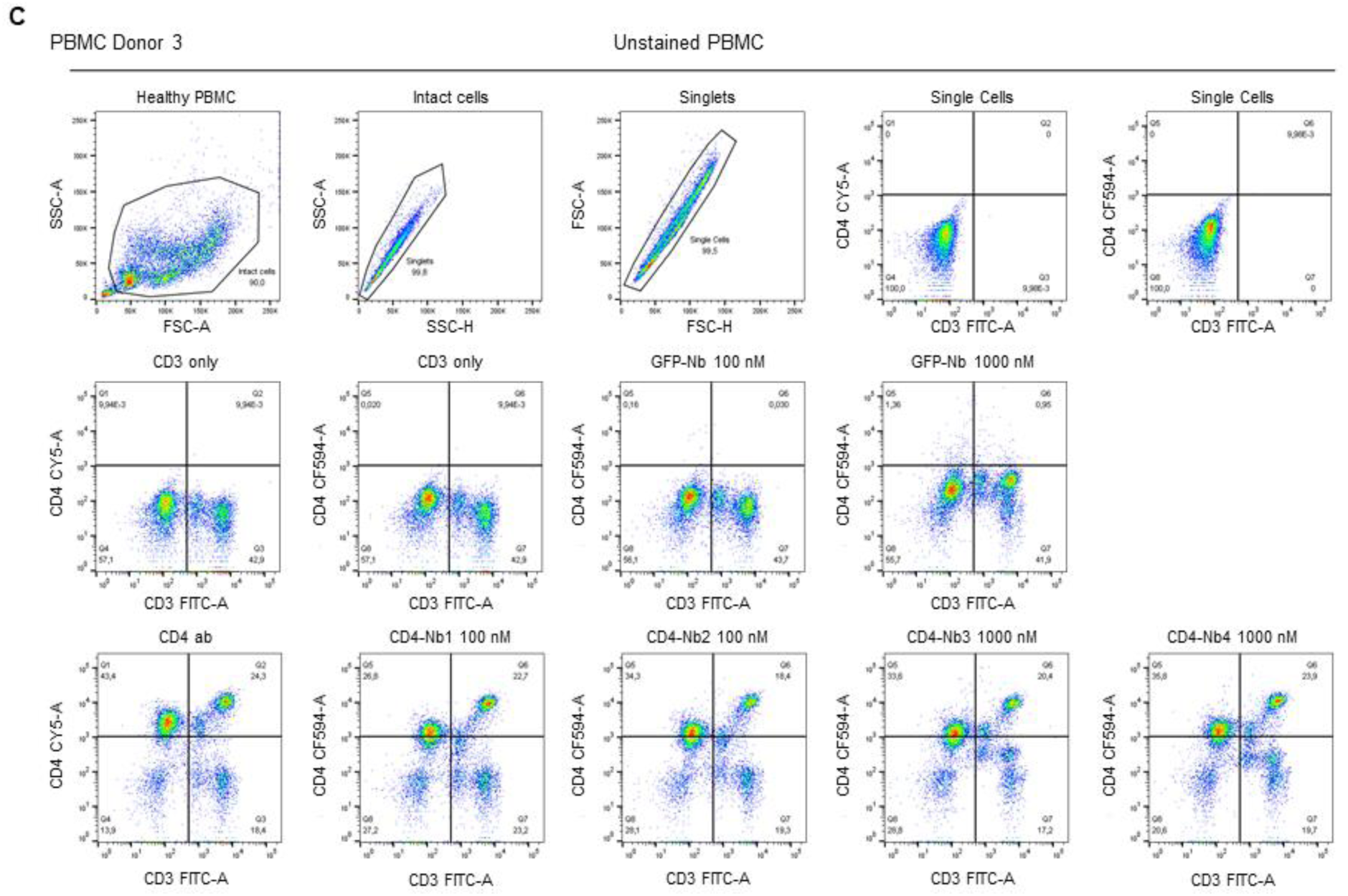
Binding of CD4-Nbs to CD4^+^ cells present in human PBMCs. Top row shows gating strategy for flow cytometry analysis of CD4^+^CD3^+^ double-positive human PBMCs. Middle and bottom row shows final gating step and quantification of these cells for donor 1 (**A**), donor 2 (**B**), and donor 3 (**C**) stained with an anti-CD4 antibody (CD4 ab), anti-GFP control Nb (GFP-Nb), or CD4-Nb1 - CD4-Nb4 at indicated concentrations.

**Supplementary Figure 7.**
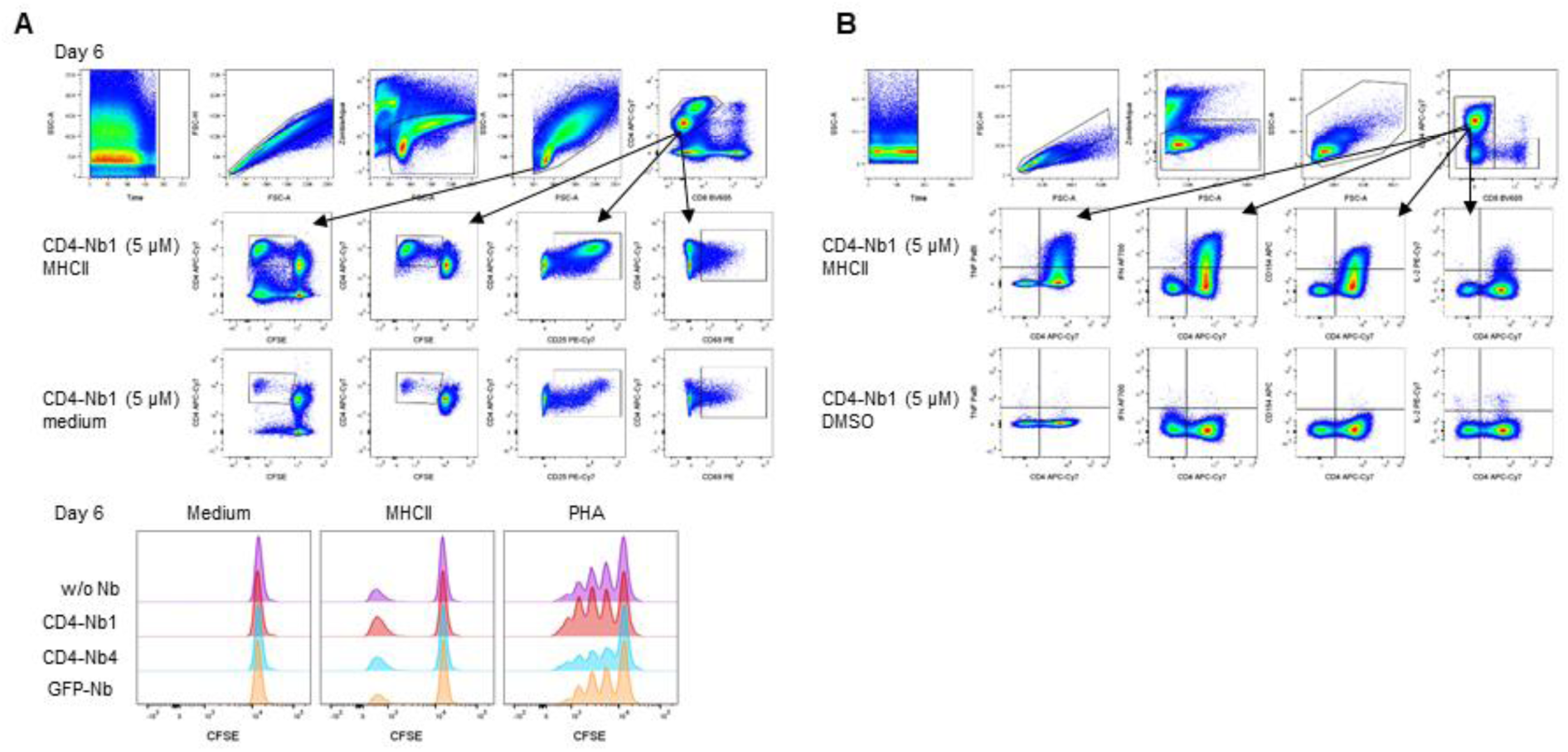
Determination of the effect of CD4-Nbs on CD4^+^ T cells. (**A**) Gating strategy for analysis of proliferation and activation of CD4^+^ cells after stimulation. Top row from left to right: Time gate, single cells, live cells, lymphocytes, CD4^+^ cells. Middle and bottom rows: gates were place on proliferating CFSE-low/negative CD4^+^ cells (left), CD25^+^CD4^+^ cells (middle) and CD69^+^CD4^+^ cells (right). Histogram overlay shows the number of divisions as CFSE labeling within CD4^+^ cells. Shown is one representative example (donor 2) on day 6. (**B**) Gating strategy (donor 3) for analysis of activation marker and cytokine expression of CD4^+^ cells in intracellular staining after 12 days of culture and 14 h restimulation. Top row from left to right: Time gate, single cells, live cells, lymphocytes, CD4^+^ cells, CD4/CD8 staining (gating on CD8^neg^ cells). Middle and bottom rows show the expression of TNF, IFN-γ, CD154 and IL-2 after restimulation with MHC-class II peptides or control DMSO/water.

**Supplementary Figure 8.**
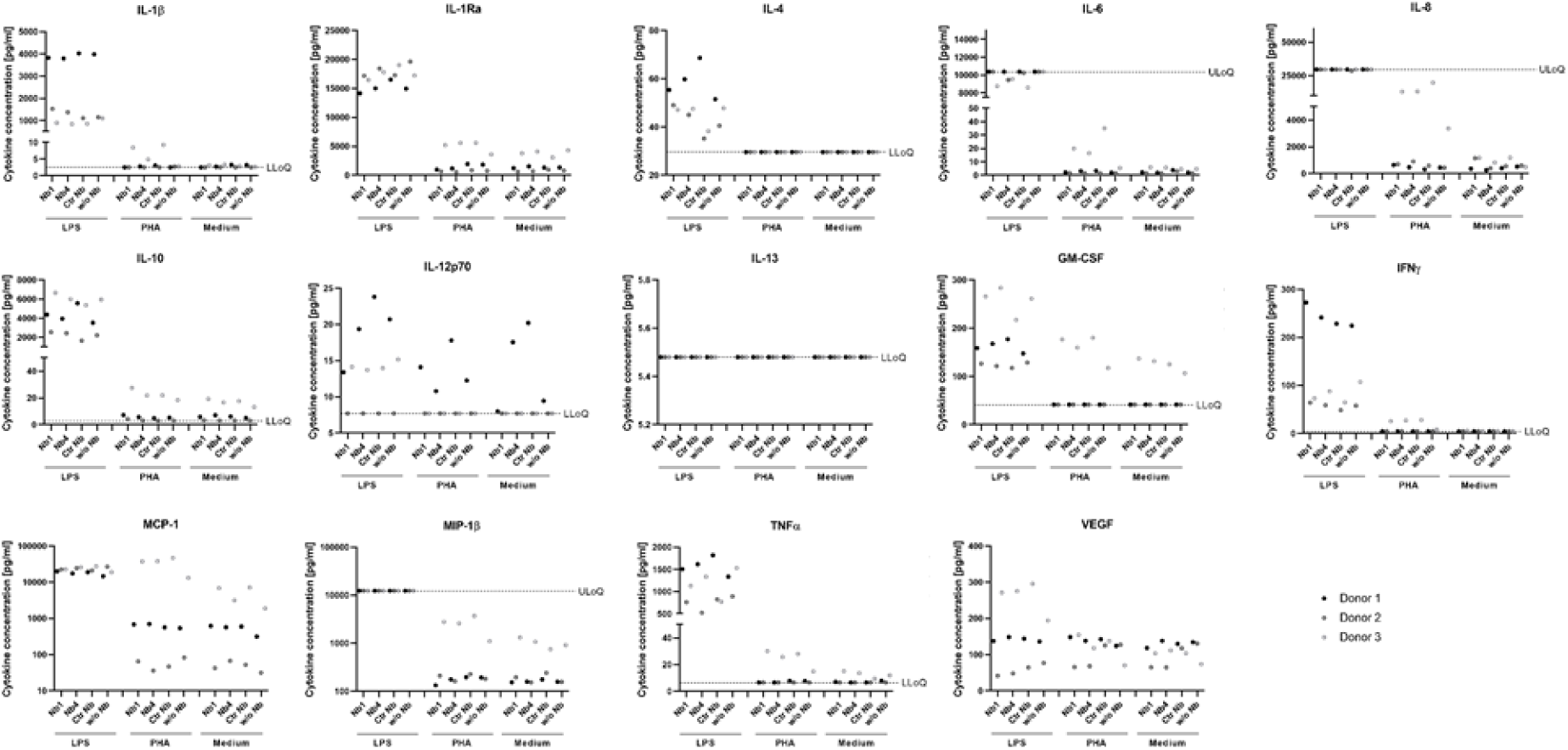
Determination of cytokines secreted from whole blood samples of donors after treatment with CD4-Nbs. Blood samples of three donors were incubated with 5µM CD4-Nb1, CD4-Nb4, GFP-Nb (control) or w/o Nb and stimulated with lipopolysaccharide (LPS), phytohaemagglutinin (PHA) or medium only as control. Secreted cytokines (listed in table S2) were measured and quantified using an in-house developed microsphere-based (Luminex) multiplex sandwich immunoassay. Results of one biological experiment are shown as colored dots indicating measured cytokine levels of one individual.

**Supplementary Figure 9.**
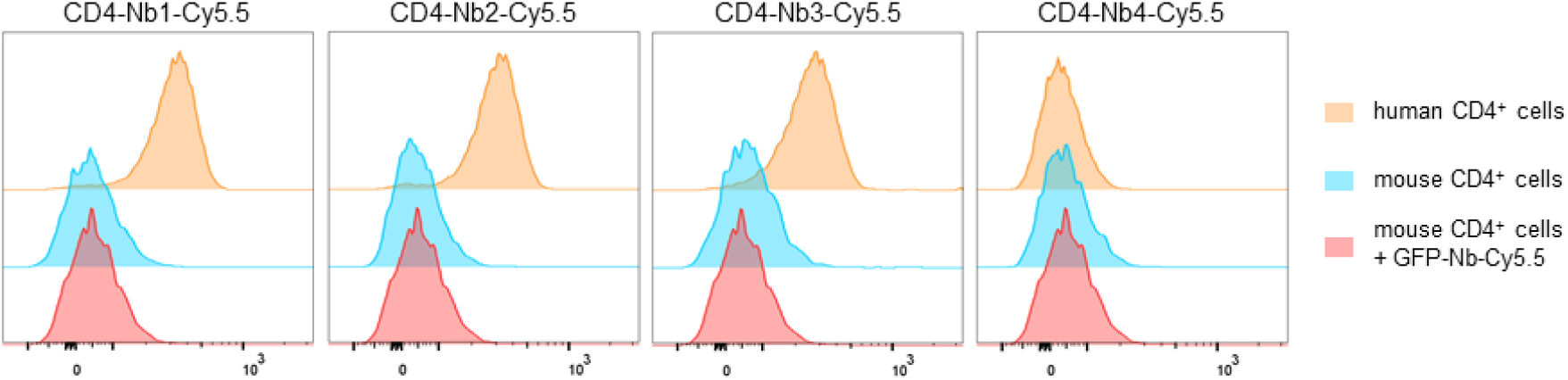
Cross-species reactivity testing of Cy5.5-labeled CD4-Nbs. Flow cytometry of human and mouse CD4^+^ cells stained with CD4-Nbs-Cy5.5 or GFP-Nb-Cy5.5. Binding to human CD4 was confirmed for CD4-Nb1, CD4-Nb2 and CD4-Nb3. CD4-Nb4 did not show staining at this concentration (0.75 µg/ml, ∼49 nM). None of the tested CD4-Nbs stained murine CD4.

**Supplementary Figure 10.**
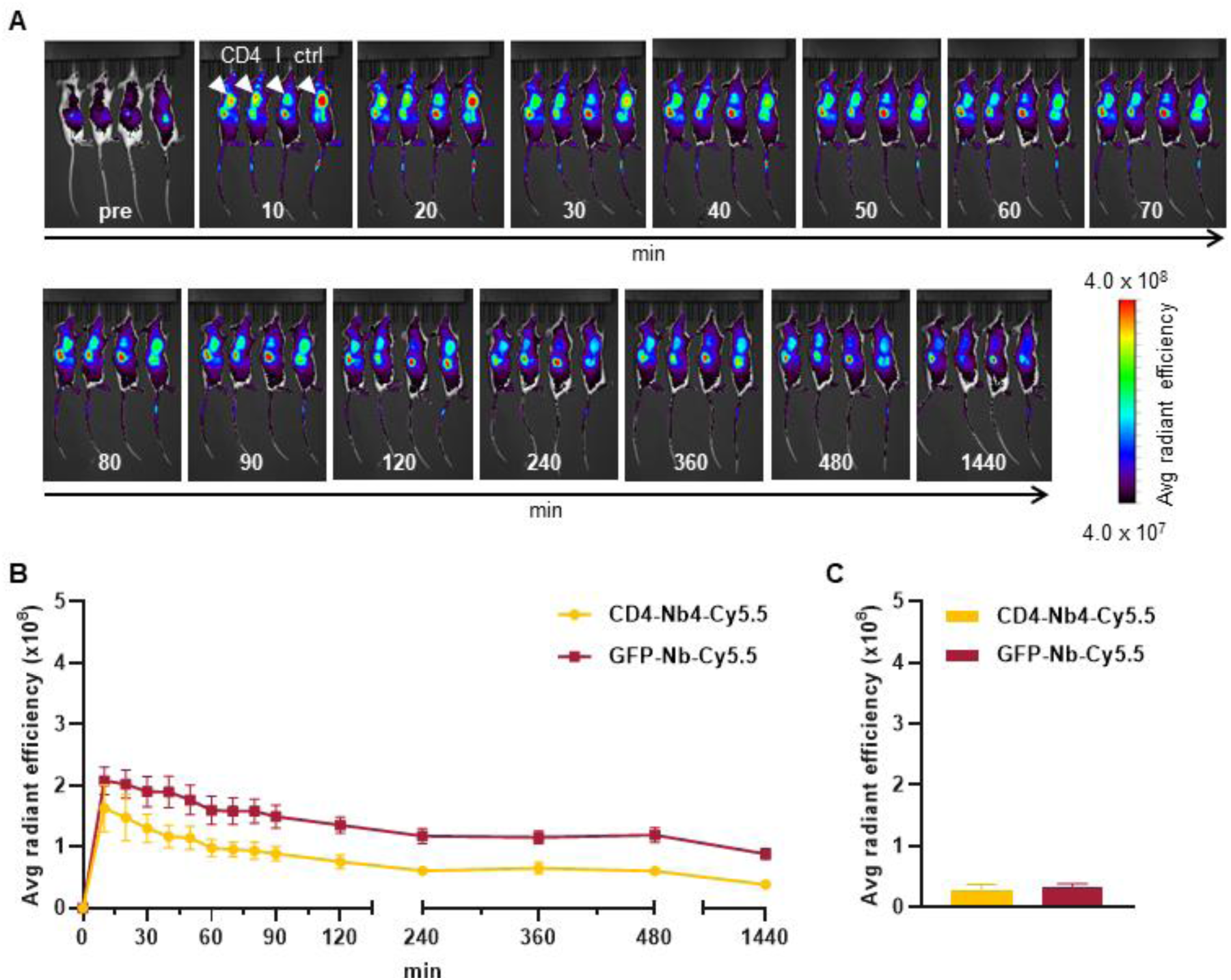
*In vivo* optical imaging (OI) of low-affinity binding CD4-Nb4-Cy5.5. 5 µg of CD4-Nb4-Cy5.5-or GFP-Nb-Cy5.5 were administered *i.v.* to *subcutaneously* human CD4^+^ HPB-ALL-bearing NSG mice and tumor bio distribution was monitored by repetitive OI measurements over the course of 24 h. (**A**) Representative images of each measurement time point of 4 mice injected either with CD4-Nb4-Cy5.5 (left, CD4) or GFP-Nb-Cy5.5 (right, ctrl). White arrows indicate the tumor localization at the right upper flank. (**B**) Quantification of the fluorescence signal from the tumors (n = 4 per group, arithmetic mean of the average radiant efficiency ± SEM). (**C**) After the last imaging time point, tumors were explanted for *ex vivo* OI, demonstrating similar accumulation of CD4-Nb4-Cy5.5 and GFP-Nb-Cy5.5 (n = 2 per group, arithmetic mean ± SEM)

**Supplementary Figure 11.**
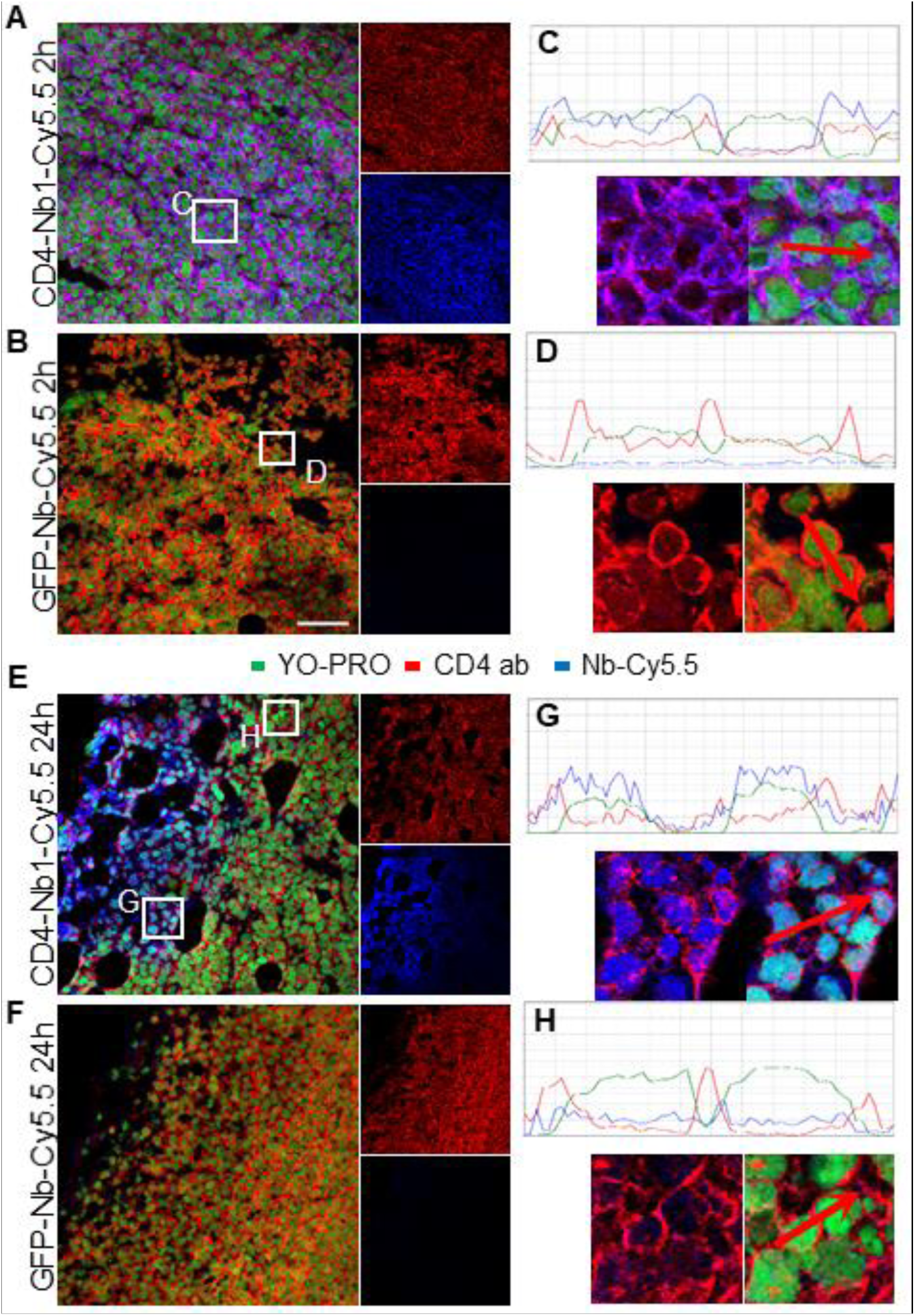
Immunofluorescence staining of e*x vivo* HPB-ALL tumors. Cryosections from *ex vivo* HPB-ALL tumors were imaged for bound Nb (blue) from the systemic injection in xenografted mice and co-stained with CD4-specific antibody (red) and nuclear YO-PRO staining (green). Image overlay of CD4-Nb1 (**A**) or control GFP-Nb (**B**) with CD4 antibody fluorescence after 2 hours of Nb injection. (**C**, **D**) Line-scan quantification of indicated image sections from A and B. (**E**, **F**) same as A and B after 24 h of systemic Nb injection. (**G**, **H**) Line-scan analysis of indicated image sections from E of strongly stained area (G) or weakly stained area (H).

**Supplementary Figure 12.**
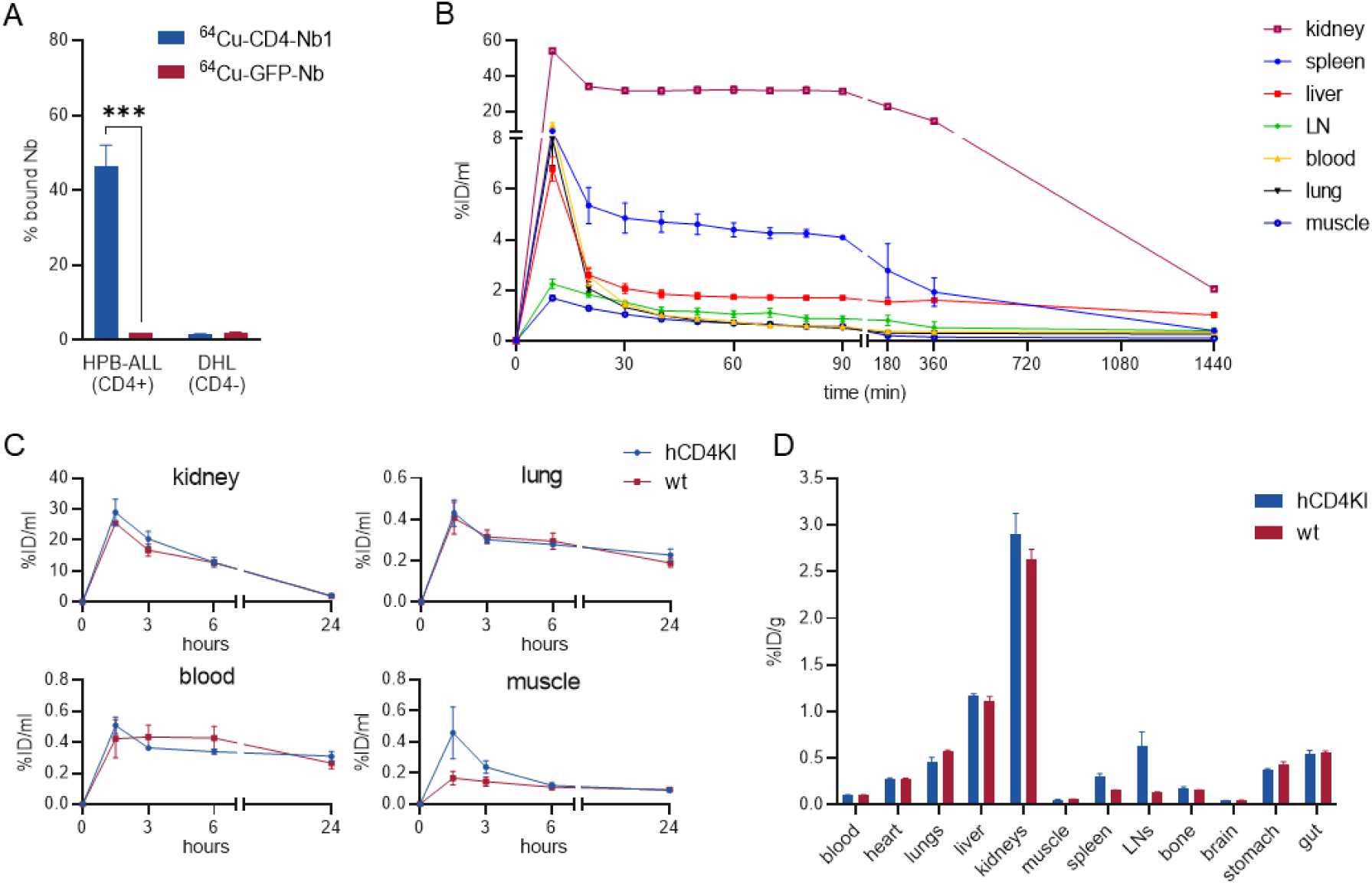
^64^Cu-CD4-Nb1 specifically accumulates in CD4^+^ T cell-rich organs. (**A**) *in vitro* binding of ^64^Cu-CD4-Nb1 or ^64^Cu-GFP-Nb to excess of CD4^+^ HPB-ALL or CD4^-^ DHL control cells analyzed by γ-counting (triplicates, arithmetic mean± SD, unpaired t-test, (***) p<0.001)). (**B**) Dynamic *in vivo* biodistribution of ^64^Cu-CD4-Nb1 in 2 hCD4KI mice by PET/MR. (**C**) Dynamic uptake quantification of ^64^Cu-CD4-Nb1 in non-T cell rich organs over 24 h (n = 3 per group). (**D**) *Ex vivo* organ biodistribution analyzed by γ-counting.

## Supplementary Methods

**Table S1:**
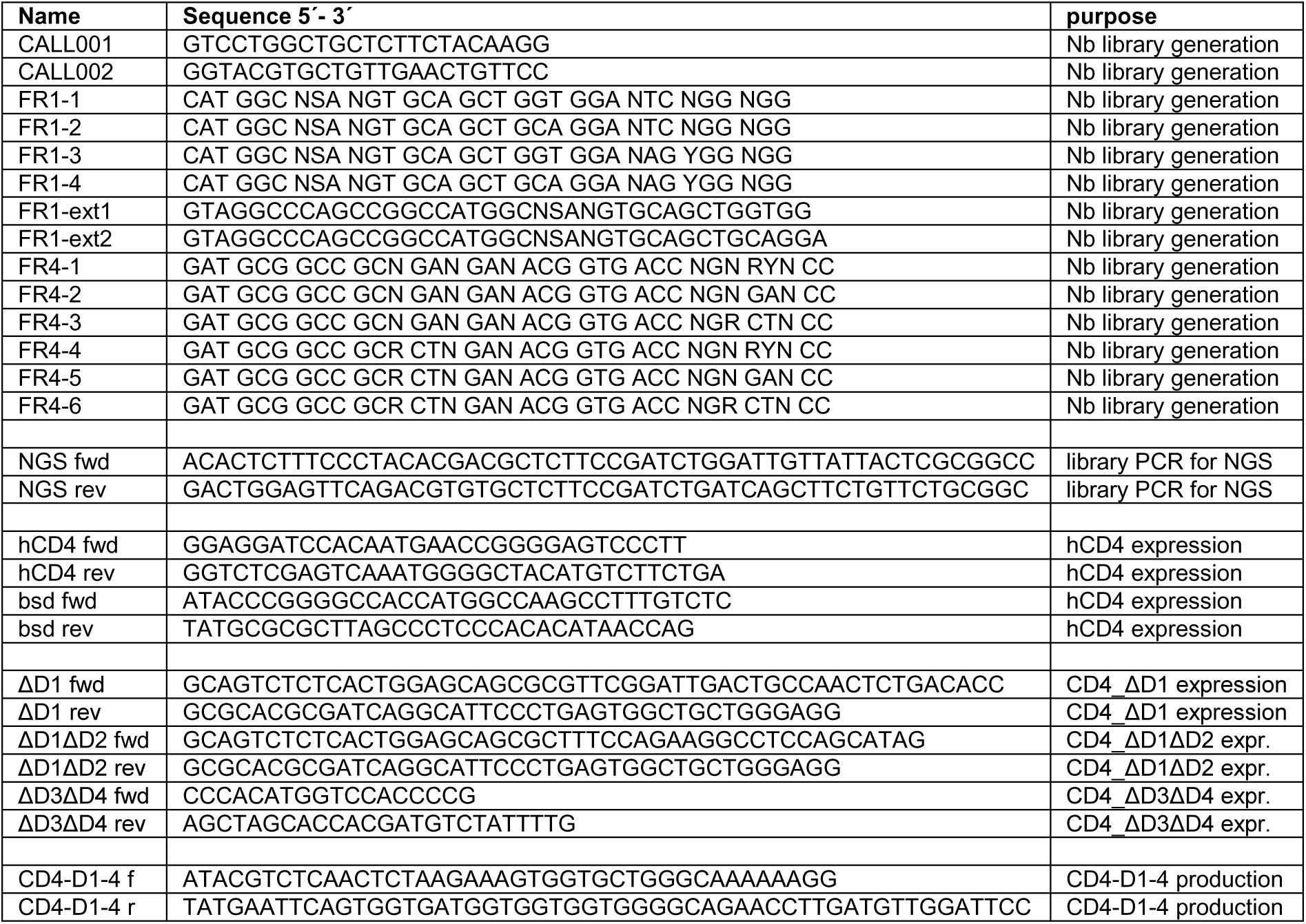
primers used in this study

**Table S2:** cytokines analyzed in this study

| **Cytokine** | **indicative for** |
| --- | --- |
| Interleukin 1 $\beta$ (IL-1β) | proinflammatory |
| Interleukin 1 receptor antagonist (IL-1RA) | antiinflammatory |
| Interleukin 4 (IL-4) | antiinflammatory |
| Interleukin 6 (IL-6) | pro-/antiinflammatory |
| Interleukin 8 (IL-8) | proinflammatory |
| Interleukin 10 (IL-10) | antiinflammatory |
| Interleukin 12 p70 (IL-12p70) | proinflammatory |
| Interleukin 13 (IL-13) | antiinflammatory |
| Granulocyte-macrophage colony-stimulating factor (GM-CSF) | pro-/antiinflammatory |
| Interferon $\gamma$ (IFNγ) | proinflammatory |
| Monocyte chemoattractant protein 1 (MCP-1) | proinflammatory |
| Macrophage inflammatory protein 1 $\beta$ (MIP-1 $\beta$) | proinflammatory |
| Tumor necrosis factor $\alpha$ (TNF-α) | proinflammatory |
| Vascular endothelial growth factor (VEGF) | wound healing factor |

**Table S3:**
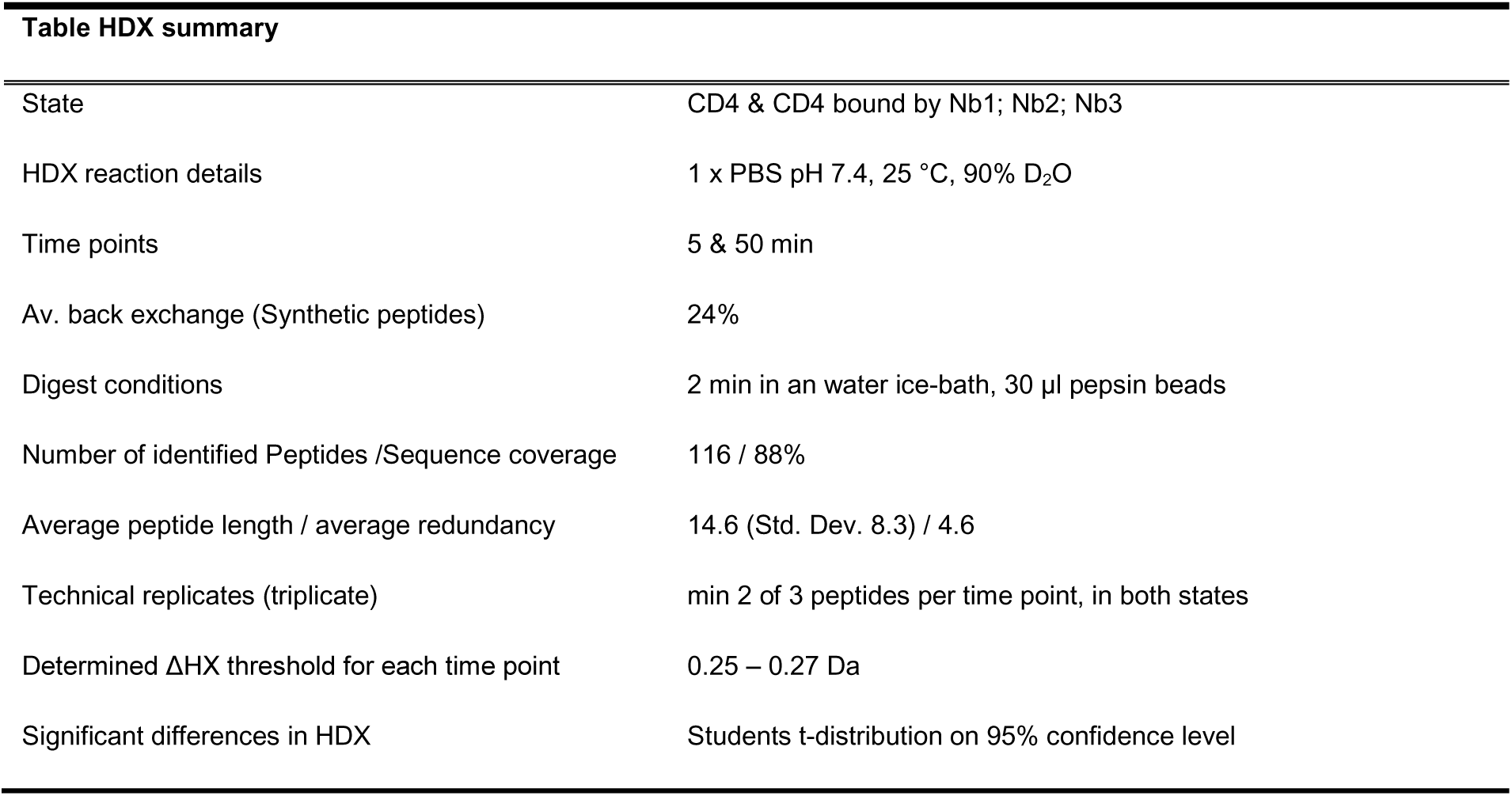
HDX-MS Summary

**Table HDX summary**
|  |  |
| --- | --- |
| State | CD4 & CD4 bound by Nb1; Nb2; Nb3 |
| HDX reaction details | 1 x PBS pH 7.4, 25 °C, 90% D<sub>2</sub>O |
| Time points | 5 & 50 min |
| Av. back exchange (Synthetic peptides) | 24% |
| Digest conditions | 2 min in an water ice-bath, 30 µl pepsin beads |
| Number of identified Peptides /Sequence coverage | 116 / 88% |
| Average peptide length / average redundancy | 14.6 (Std. Dev. 8.3) / 4.6 |
| Technical replicates (triplicate) | min 2 of 3 peptides per time point, in both states |
| Determined ΔHX threshold for each time point | 0.25 – 0.27 Da |
| Significant differences in HDX | Students t-distribution on 95% confidence level |

